# *EnhancerDetector:* Enhancer Discovery from Human to Fly via Interpretable Deep Learning

**DOI:** 10.1101/2025.05.28.656532

**Authors:** Luis M. Solis, Geyenna Sterling-Lentsch, Marc S. Halfon, Hani Z. Girgis

## Abstract

**Background:** Deciphering how enhancers encode regulatory information in DNA remains a central genomics challenge, as sequencing outpaces functional annotation. A key question is whether enhancers possess an intrinsic, sequence-based “enhancerness” distinguishing them from other regions, independent of species, cell type, or assay. Confirming its existence and learnability is both biologically fundamental and essential for scalable genome annotation.

**Results:** We introduce *EnhancerDetector*, a convolutional neural network-based framework for cross-species enhancer prediction that combines high accuracy with biological interpretability. Trained on human data, *EnhancerDetector* achieves strong performance across human, mouse, and fly datasets, consistently outperforming existing methods in precision and F1. It generalizes to datasets generated using diverse experimental assays. Unlike chromatin feature-based predictors requiring complex post hoc thresholding, *EnhancerDetector* directly outputs enhancer probability scores from short sequence windows, simplifying enhancer discovery workflows. An ensemble strategy further improves prediction reliability by reducing false positives. *EnhancerDetector* supports fine-tuning on new species and retains strong performance even when adapted with as few as 20,000 enhancer sequences, making it ideal for newly sequenced genomes with limited experimental data. For interpretability and visualization, we apply class activation maps to identify sequence regions predictive of enhancer activity. Experimental validation in transgenic flies confirms the predictive power of *EnhancerDetector* : five of six tested candidates drove reporter expression, and four exhibited expression patterns supported by prior literature. These analyses highlight distinct sequence and contextual features that confer what we term “enhancerness:” enhancer sequences possess a characteristic, identifiable signature.

**Conclusions:** *EnhancerDetector* provides a sequence-based and interpretable framework for enhancer discovery across species and experimental contexts. Our results support the presence of recurring enhancer-associated sequence features that can be learned from DNA sequence and transferred across genomes. By combining cross-species prediction, fine-tuning, model interpretation, and experimental validation, *EnhancerDetector* offers a practical approach for prioritizing candidate enhancers in well-studied genomes. These features position *EnhancerDetector* as a useful first-stage annotation tool for identifying putative enhancers in newly sequenced genomes with limited experimental data.

## Background

Enhancers are regulatory elements that play a critical role in the transcriptional activation of genes. These non-coding DNA sequences can be located upstream, downstream, or even within the target gene itself, and their ability to function across long genomic distances is facilitated by the three-dimensional architecture of a genome [1, 2]. Enhancers play a crucial role in controlling gene activity that underlies lineage commitment and organismal formation [3]. Moreover, enhancer dysfunction is increasingly recognized as a contributor to various medical issues, collectively termed “enhanceropathies” [1]. Thus, accurate identification of enhancers is essential for understanding gene regulation in both physiology and pathology.

Despite their biological significance, identifying enhancers remains a considerable challenge. Enhancers exhibit high locational variability and are often found within extensive regions of non-coding DNA, making them difficult to detect [2]. This challenge is compounded by the cell-type-specific nature of enhancer activity: an enhancer active in one cellular context may be inactive in another [2]. Experimental methods—reviewed by Suryamohan and Halfon [4]—such as ChIP-seq, DNase-seq, ATAC-seq, FAIRE-seq, Hi-C, classical reporter assays, high-throughput reporter screens, CRISPR-based perturbation techniques, and eRNA-detection assays have advanced enhancer discovery [5–19]. However, each captures only a fraction of the enhancer landscape, requires substantial experimental resources, and is typically applied to a narrow set of species, tissues, and developmental stages. As a result, even in well-studied organisms, enhancer annotations remain incomplete.

The gap is even more striking in the rapidly expanding collection of newly sequenced genomes. Large-scale initiatives such as the Earth BioGenome Project [20], the Vertebrate Genomes Project [21], the European Reference Genome Atlas [22], and global marine surveys such as Tara Oceans [23] are generating reference genomes and environmental sequence resources far faster than functional genomic data can be annotated. Because the vast majority of these species lack ATAC-seq, ChIP-seq, or reporter assay data, their regulatory landscapes—especially enhancers—are effectively unannotated. Thus there is a critical need for scalable approaches that can identify putative enhancers from DNA sequence alone, enabling first-pass functional annotation across thousands of species.

A key question, therefore, is whether enhancers possess generalizable sequence-level properties that can be detected independently of cell type or species [24]. Biologically, enhancers are enriched for dense clusters of transcription factor binding sites (TFBSs) [25, 26]. These binding sites are often degenerate and exhibit substantial similarity across transcription factor families—for example, the CANNTG core for bHLH factors, TAAT for homeobox proteins, and GGA/T for ETS proteins [27]. Although the precise combinations and arrangements of motifs that drive enhancer activity are cell-type specific, the enrichment of such degenerate motif clusters is a shared and fundamental property of enhancers [25]. These observations support the hypothesis that enhancers are defined by intrinsic, recurring sequence characteristics—suggestive of a general enhancer-defining signature—that distinguish them from other genomic regions, even when their regulatory outputs are context dependent [28, 29].

We therefore hypothesize that enhancers possess an intrinsic and conserved sequence signature—here termed “enhancerness”—that distinguishes enhancer DNA from non-enhancer DNA. This signature arises from the recurrent presence and density of related transcription factor motif families, their short-range organization, and supportive broader sequence context [30]. Under this hypothesis, a model trained on a diverse collection of enhancers from multiple cell types should be able to learn enhancer identity even when the specific transcriptional codes governing activity differ.

Existing computational methods have provided important advances in enhancer prediction but also exhibit several limitations. Early approaches relied on traditional machine learning models trained on manually designed sequence features (e.g., k-mer frequencies, GC content, or motif scores), exemplified by tools such as CrmMiner [31], iEnhancer-2L [32], EnhancerPred [33], iEnhancer-EL [34], and SCRMshaw [35]. More recent methods have adopted deep learning strategies, including convolutional, recurrent, generative, and attention-based architectures trained directly on DNA sequence [36–39]. In parallel, widely used models such as gkm-SVM and LS-GKM [40, 41], DeepSEA [42], deltaSVM [43], and Enformer [44] have demonstrated strong performance and broad utility in regulatory genomics; however, these approaches are typically optimized for predicting activity in specific cellular contexts rather than for identifying generalizable enhancer identity.

Despite these advances, many existing approaches rely on chromatin marks or handcrafted features, limiting their applicability to species and tissues with rich functional genomic datasets, while others require post hoc thresholding or show limited generalization across species, assays, or experimental contexts. Sequence-only deep learning methods offer a promising alternative, but many were not explicitly designed to capture generalizable enhancer identity or to provide interpretable insights into the sequence features underlying enhancer function [45].

To address these challenges and to directly test whether enhancers share a generalizable, sequence-encoded identity, we developed *EnhancerDetector*, a convolutional neural network-based framework for identifying enhancers from DNA sequence alone. *EnhancerDetector* is trained on curated enhancer datasets from human and is designed to generalize across species, datasets, and experimental assays. By learning sequence-level properties that distinguish enhancers from non-enhancers, the model provides a scalable solution for predicting putative enhancers in genomes lacking functional genomic data. Moreover, by incorporating class activation maps, *EnhancerDetector* yields interpretable insights into the sequence features associated with enhancer identity. Experimental validation in transgenic flies demonstrates that these predictions correspond to biologically active enhancers.

In this study, we show that enhancers possess a detectable sequence signature and that this signature can be learned in a manner that generalizes across species—a property with significant implications for genome annotation. Our results provide evidence for the existence of enhancerness as a generalizable feature of enhancer sequences, and they introduce a practical framework for enhancer discovery at a genomic scale.

## 1 Methods

### 1.1 Overview

*EnhancerDetector* is a flexible enhancer prediction framework that uses deep learning to classify genomic sequences as enhancers or non-enhancers. Given a sequence in FASTA format, the tool outputs a confidence score representing the probability that the input sequence functions as an enhancer. *EnhancerDetector* is centered on a convolutional neural network, with an optional ensemble mode to boost prediction confidence. These models are trained on curated enhancer datasets from both human and fruit fly genomes, and support fine-tuning for application to new species using limited enhancer data. *EnhancerDetector* also employs class activation maps, an explainable AI technique that highlights the most influential regions within input sequences. An overview of the architecture is shown in Figure 1.

**Figure 1:**
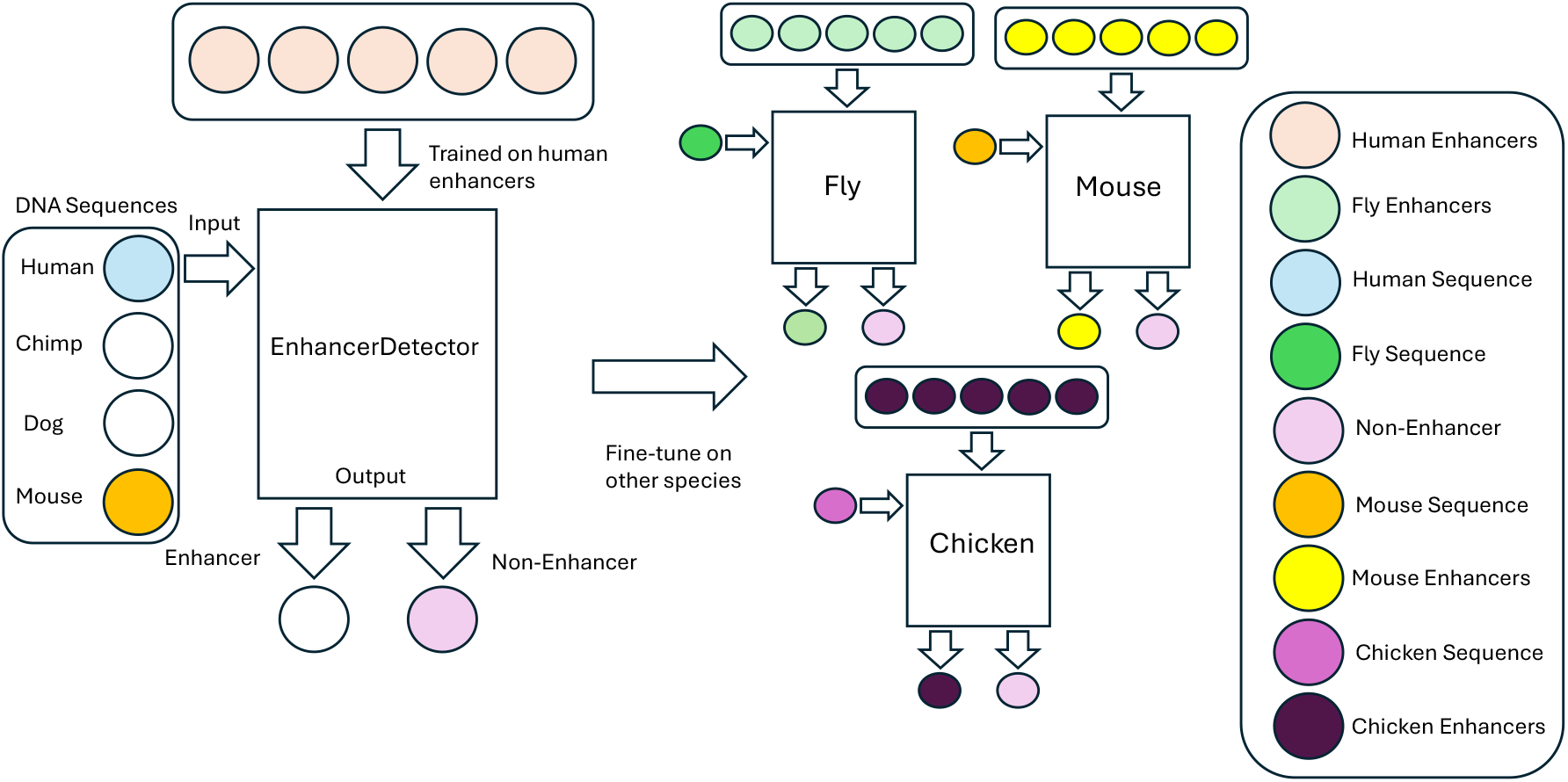
General Architecture and Cross-Species Adaptability of *EnhancerDetector*. The left panel depicts the base model trained on the human CATlas dataset, which can be directly applied to closely related vertebrate species with minimal performance degradation. The right panel highlights *EnhancerDetector* ‘s support for fine-tuning, enabling the human-trained model to be adapted for more distantly related species. For illustrative purposes, fine-tuning is shown for fruit fly, mouse, and chicken, though the approach is broadly applicable to any species with sufficient enhancer data.

### 1.2 Enhancer Dataset

For the human classifier, we utilized a comprehensive set of putative enhancer sequences curated from the Cis-element Atlas (CATlas), which we will henceforth refer to as the CATlas dataset. This dataset was generated through the application of single-nucleus open chromatin accessibility sequencing (snATAC-seq) across 222 distinct cell types derived from both adult and fetal human tissues. The identified regions of open chromatin should encompass a diverse repertoire of regulatory elements, including transcriptional enhancers. The sequences within the CATlas dataset are uniformly 400 base pairs (bp) in length. For the *Drosophila melanogaster* classifier, we integrated distal DNase I hypersensitive sites identified by the Stamatoyannopoulos Laboratory [46], distal scATAC-seq regions from the Furlong Laboratory [47], and experimentally validated *cis*-regulatory modules (CRMs) sourced from REDfly [48].

### 1.3 Preprocessing

To ensure the high quality of our enhancer datasets, we implemented a stringent preprocessing pipeline to exclude potentially confounding genomic features, namely promoters, coding regions (exons), and insulators, from the human, mouse, and fly datasets. For the human data, we leveraged RefSeq and GENCODE annotations to delineate transcription start sites and the genomic coordinates of coding exons. For the mouse data, promoters and exons were obtained from Ref-Seq and GENCODE VM25 annotations via the UCSC Genome Table Browser. For the fly data, we employed the RefSeq and Ensembl databases to obtain analogous information. We defined a promoter region as a 1000 bp region centered on each annotated transcription start site. Human and mouse insulator regions were retrieved from the ENCODE Project via the UCSC Genome Table Browser, specifically selecting regions characterized by the presence of CTCF binding sites. For the fly, we incorporated putative insulator regions derived from ChIP-based binding profiles of known insulator-associated proteins from a previously published study [49]. All instances of overlap between these potentially confounding regions and our initial enhancer datasets were systematically removed using BEDTools [50]. For the mouse ENCODE dataset, regions shorter than 400 bp after preprocessing were symmetrically expanded to 400 bp to match the input length expected by the human-trained network. Finally, to ensure uniformity in genomic coordinates for the fly data, all sequences were mapped to the DM6 genome assembly using the LiftOver tool [51].

### 1.4 Control Datasets

To rigorously evaluate our model’s ability to discriminate true enhancers from non-functional genomic regions, we generated five distinct control datasets representing negative examples for each species. Four of these datasets were specifically designed to control for potential confounding factors: sequence length, repetitive element content, and guanine-cytosine (GC) content. Based on these variables, we constructed the following datasets:

- Length Repeat (LR): Sequences in this control set were sampled to match the length distribution of the enhancer dataset, allowing for the presence of repetitive elements.
- Length No-Repeat (LNR): This dataset comprises sequences sampled to match the length distribution of the enhancer dataset, with the exclusion of repetitive elements.
- Length GC-Repeat (LGR): Sequences in this control set were sampled to match both the length and GC content distributions of the enhancer dataset, while permitting the inclusion of repetitive elements.
- Length GC-No-Repeat (LGNR): This dataset contains sequences sampled to match both the length and GC content distributions of the enhancer dataset, with the exclusion of repetitive elements.

In addition to these controlled datasets, we generated a fifth control set consisting of shuffled enhancer sequences. For each enhancer in our primary datasets, we generated shuffled sequences by randomly permuting its constituent k-mers (subsequences of length *k*), where *k* ranged from 1 to 6. This control aims to assess the model’s capacity to distinguish true enhancers from sequences that merely share the same k-mer composition. Repetitive elements in the human, mouse, and *Drosophila melanogaster* genomes were identified using Red [52]. Control regions for the three species were selected from their respective genomes, ensuring no overlap with the enhancer datasets. When generating control sequences for the human and mouse datasets, lengths were precisely matched to 400 bp. For the *Drosophila melanogaster* dataset, control sequence lengths were randomly chosen from a uniform distribution between 460 and 500 bp for each corresponding enhancer, providing a broader range for scanning. A tolerance of 0.03 was permitted for the GC content of the selected control sequences in the human, mouse, and *Drosophila melanogaster* datasets.

### 1.5 Data Partitioning

Our enhancer and control datasets were split into training (70%), validation (20%), and testing (10%) datasets. Given the requirement for deep neural networks to be effectively trained with substantial amounts of data, we allocated 70% of our dataset for training purposes. An additional 20% of the data was designated as the validation dataset to detect and mitigate overfitting—the phenomenon where a model learns the training data too well but fails to generalize to new, unseen data (i.e., memorizing instead of learning). The remaining 10% of the data served as the testing dataset to ensure a comprehensive evaluation on previously unseen data. The data from both enhancers and controls in the human dataset were randomly allocated across these partitions. This random split ensures that each partition contains a representative mix of both enhancers and non-enhancers. However, for the fly dataset, a different approach was required. Since the enhancers for the fly were sampled from four different experiments there is potential overlap between the enhancers. If enhancers were split randomly, overlapping enhancers could appear across two or three partitions, leading to data leakage. As a remedy, sequences were sorted by chromosome and start position. After sorting, the data were split sequentially, ensuring that enhancers with overlapping genomic coordinates were assigned to the same partition, thus eliminating potential data leakage between training, validation, and testing sets.

### 1.6 GC Distribution

After partitioning the data, we ensured that the enhancer and control sequences were matched in terms of length and GC content. This prevented the model from oversimplifying the classification task by relying on differences in these basic sequence properties. As shown in Figure 2, the GC content distributions for both the fly and human datasets were well matched between enhancers and controls. To quantify the similarity between the enhancer and control datasets, we used Kullback-Leibler (KL) divergence to compare the distributions of sequence length and GC content. A KL divergence of zero indicates identical distributions, with lower values reflecting greater similarity. For the fly datasets, we observed KL divergence values ranging from 0.1173 to 0.2216 for training and from 0.1272 to 0.2267 for validation. For the human datasets, values ranged from 0.0068 to 0.2233 for training and from 0.0066 to 0.2236 for validation, indicating strong similarity in both species. Shuffled control datasets yielded KL divergence values of 0.0 because they were generated directly from the enhancer sequences.

**Figure 2:**
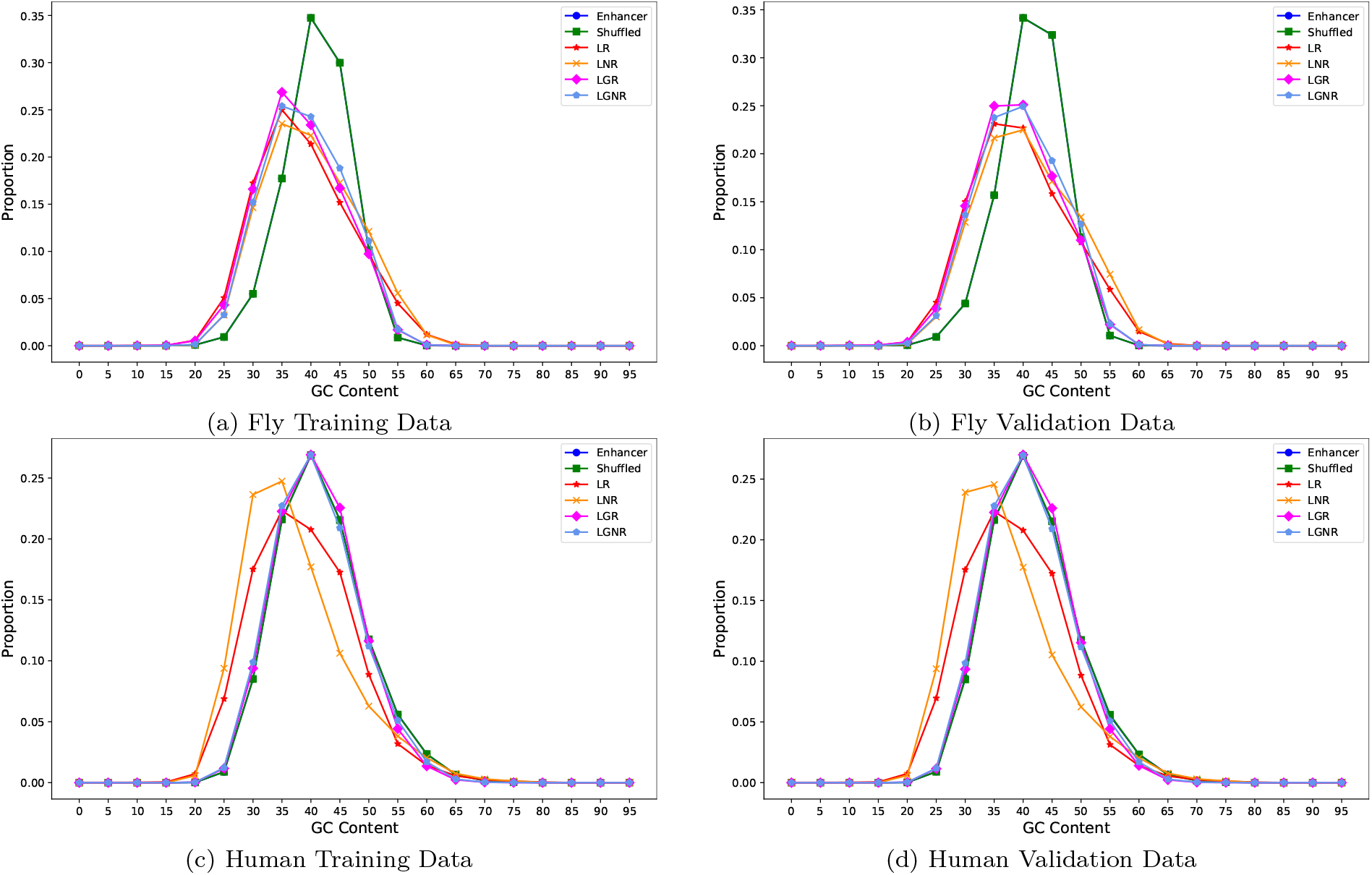
Comparison of GC content in fly and human enhancer training and validation datasets. This figure illustrates the similarity in GC content distributions between control and enhancer datasets for both training and validation splits in human and fly. All four panels show closely aligned GC content distributions, demonstrating that the control sequences are well matched to the enhancers in terms of GC content. Legend abbreviations: Enhancer is the true enhancer sequences. LR is control sequences matched for length with repeats allowed. LNR is length-matched controls with repeats removed. LGR is controls matched for length and GC content with repeats allowed. LGNR is length- and GC-matched controls with repeats removed. Shuffled represents enhancer-derived sequences with permuted k-mer composition.

### 1.7 Data Indexing

To enable the neural network to learn a representation of each nucleotide, we assigned a unique integer index to each nucleotide. The specific values are unimportant to the network, as long as they are positive and unique. We modified the input sequences by assigning an integer to each nucleotide encountered in the genomic data. For example, the nucleotide A is assigned the index 2, with 0 and 1 reserved for padding and unknown bases, respectively. T is assigned the index 3, and so on, until every nucleotide has been mapped to a numerical value. Thus, a sequence such as ATAT is encoded as [2, 3, 2, 3], which can be used as input to the network.

### 1.8 Data Augmentation

To train the network effectively, we increased the amount of training data through a data augmentation strategy. We generated additional data by creating reverse complement counterparts of the original sequences and using them as input. This approach is valid because forward and reverse complement sequences are biologically equivalent. As a result, the model was exposed to more diverse training examples without requiring additional data collection. During training, each input sequence had a 50% chance of being replaced with its reverse complement. These reverse complements were indexed into numerical representations using the same scheme as the forward sequences. We applied an additional augmentation technique when preparing the fly dataset. To ensure that all sequences fell within a uniform length range (460– 500 bp), we expanded or slightly trimmed sequences as needed. This was important as we planned to scan long sequences by dividing them into segments within this range. The procedure was the same for both expansion and trimming: we determined the midpoint of each sequence, then generated five sequence copies of randomly sampled lengths between 460 and 500 bp, all centered on the midpoint.

### 1.9 Convolutional Neural Network

For *EnhancerDetector*, the primary model used is a convolutional neural network. Convolutional neural networks are well-suited for processing data with a structured arrangement, such as DNA sequences, and are effective at identifying and learning hierarchical patterns. Our network (Figure 3) begins with an embedding layer, which learns numerical representations for nucleotides. The output of the embedding layer is passed to four custom convolutional blocks, each consisting of two one-dimensional convolutional layers followed by a batch normalization layer. The output of the final block is flattened and passed through two dense layers, with a batch normalization layer in between. The network ends with a single-neuron dense layer that outputs a probability score between 0 and 1, representing the likelihood that a given sequence is an enhancer. Sequences with scores greater than or equal to 0.5 are classified as enhancers, while those below 0.5 are classified as non-enhancers. We optimized the following hyperparameters:

**Figure 3:**
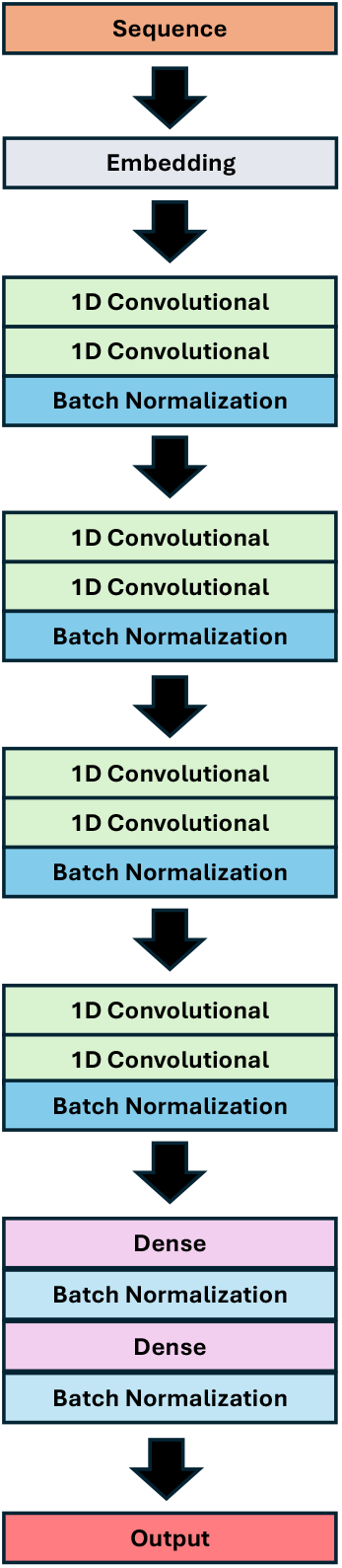
An overview of the *EnhancerDetector* architecture. The network takes a 400-bp-long DNA sequence as input and passes it through an embedding layer, which learns a numerical representation for each nucleotide. The resulting sequence is then processed by four convolutional blocks, each consisting of two 1D convolutional layers (kernel size = 3), where the second uses a stride of 2 for down-sampling. These are followed by batch normalization and a ReLU activation. The number of filters in each block increases with depth (64, 128, 256, and 512), allowing the network to extract hierarchical features of increasing complexity. Next, the output is flattened and passed through two dense layers (each with 20 units and ReLU activation), with a batch normalization layer in between, to classify whether a given sequence is an enhancer. The final output is a value between 0 and 1, where values above 0.5 indicate enhancers.

- Number of filters: Determines how many patterns each convolutional layer learns.
- Filter size: Specifies the width of the filters, e.g., a size of 3 corresponds to patterns spanning three consecutive positions.
- Number of dense neurons: Controls the number of neurons in the dense layers used for classification.

### 1.10 Recurrent Neural Network

Another model we considered was a recurrent neural network, which is particularly well-suited for sequential data, as it can capture temporal dependencies and patterns over time. This model uses a recurrent block consisting of a Gated Recurrent Unit (GRU) layer, followed by a batch normalization layer and an activation function. The architecture is similar to that of the convolutional neural network: it begins with an embedding layer, followed by two recurrent blocks, then two dense layers with a batch normalization layer in between. For this network, we optimized the following three hyperparameters: the number of GRU neurons, the activation function used in the recurrent block, and the number of dense neurons.

### 1.11 Evaluation Metrics

We evaluated the performance of *EnhancerDetector* and related tools using the following metrics (higher values indicate better performance across all metrics):

- Accuracy: Percentage of correctly predicted samples (enhancers or non-enhancers).
- Recall (Sensitivity): Percentage of true enhancers correctly identified.
- Specificity: Percentage of true non-enhancers correctly identified.
- Precision: Proportion of predicted enhancers that are correct. This is especially important for experimental validation in transgenic flies.
- F1 and Weighted F1 Scores: The F1 score is the harmonic mean of precision and recall. The weighted F1 score also incorporates precision and recall, but precision was weighted twice as much.

### 1.12 Guarding Against Overfitting

In addition to the data augmentation techniques described earlier, we applied early stopping to prevent overfitting—a situation where the model performs well on training data but poorly on unseen data. Early stopping is a regularization method that halts training when performance on validation data stops improving.

Specifically, we monitored the weighted F1 score on the validation set after each epoch (one complete pass through the entire training dataset). Training continued as long as the score improved. If no improvement was observed for 25 epochs, training was stopped and the model with the highest validation weighted F1 score was selected.

### 1.13 Optimization

The models were evaluated using various combinations of hyperparameters to identify the optimal configuration. The following hyperparameter values were tested: number of filters (32, 40, 48, 64, and 72), filter size (3 and 5), and number of dense neurons (10, 20, and 40). The best configuration for the human enhancer classifier was a combination of 64 filters, a filter size of 3, and 20 dense neurons. Similarly, we identified two best networks for the fly enhancer classifier, which were optimized with combinations of 32 and 40 filters, a filter size of 3, and 20 dense neurons. The best models were primarily chosen based on a high weighted F1 score, due to the imbalance between confirmed enhancers and control data. Specificity was also considered to ensure the model does not incorrectly identify non-enhancers, and recall was assessed to verify the model’s ability to correctly identify enhancers.

### 1.14 Fine-Tuning

We observed that the human model was trained on a comparatively larger dataset than the fly model (8,954,140 vs. 6,473,088), allowing it to learn certain characteristics and patterns more effectively. We therefore decided to adapt our trained human classifier to the fly dataset using a fine-tuning approach. The rationale was to adjust the model to optimize its performance on a new but related dataset. To fine-tune the classifier on the fly data, we employed a strategy of freezing and unfreezing layers during training. Only the unfrozen layers were updated with the new data, enabling the model to adapt while preserving knowledge gained from the human data. This approach allows us to retain useful features learned from the human dataset while tailoring the model to the specific characteristics of the fly data.

### 1.15 Ensemble

After training the fly networks and fine-tuning the model, we created an ensemble of these three networks. An ensemble combines multiple neural network models, each independently tasked with determining whether a given genomic region is an enhancer. The ensemble aggregates the predictions from all three models: if all agree that a region is an enhancer, it is classified as such; otherwise, it is not. This approach provides a stricter criterion for identifying enhancer regions, reducing the likelihood of false positives.

### 1.16 Comparison to Related Tools

To further validate *EnhancerDetector*, we compared its performance with that of other enhancer detection tools designed for binary classification. One such tool is Enhancer-FRL [53], which predicts human enhancers of 200 bp in length using feature representation learning and multi-view sequence information. We evaluated Enhancer-FRL using two datasets: the ENCODE Human dataset (obtained from the UCSC Genome Table Browser) and the ENCODE Mouse dataset. Both datasets were preprocessed as described earlier, including the removal of promoters, exons, and insulators. For the Enhancer-FRL evaluation, we selected potential enhancers that were 200 bp or shorter. Regions shorter than 200 bp were symmetrically extended in both directions to reach exactly 200 bp in length. This same procedure was applied to mouse enhancers. Because our human network expects input sequences of 400 bp, we further extended the 200-bp enhancers to 400 bp. To ensure a fair comparison, we excluded any enhancer region that overlapped by at least 50% with our CATlas training enhancer dataset. This filtering ensured that none of the evaluated enhancers had been seen during training. For both the 200-bp and 400-bp enhancers in human and mouse datasets, we also generated matched control sequences (shuffled, LR, LNR, LGR, and LGNR). To ensure realistic and biologically relevant benchmarking, all model evaluations were conducted using testing datasets with a 1:10 enhancer-to-control ratio for both the human and mouse ENCODE datasets. This ratio reflects the natural sparsity of enhancer elements in the genome and more accurately simulates genome-wide classification scenarios. Enhancer-FRL was run using its default parameters.

Another tool we compared *EnhancerDetector* to was Enhancer-LSTMAtt [54]. This method detects 200-bp enhancers using a deep neural network that incorporates bidirectional Long Short-Term Memory (LSTM) layers and an attention mechanism. Enhancer-LSTMAtt was evaluated using the same 200-bp human and mouse ENCODE datasets, along with their corresponding control datasets. The model was executed using its default parameters.

A third tool we evaluated was the PDCNN model [55]. PDCNN is a convolutional neural network specifically designed for DNA sequence analysis. Unlike the other tools, PDCNN was trained on both human and mouse enhancers. The authors used sequences ranging from 50 bp to 300 bp for testing and recommended using 200-bp sequences for optimal performance. Accordingly, we evaluated PDCNN using the same 200-bp human and mouse enhancer datasets, along with their corresponding control datasets. PDCNN was run using its default parameters.

Another tool we assessed was gkm-SVM [40]. This sequence-based classifier uses gapped k-mer features and support vector machines to predict functional regulatory elements in genomic sequences. By allowing gaps within k-mers, gkm-SVM captures more flexible and informative sequence patterns compared to traditional k-mer methods. More specifically, we used LS-GKM [41], an optimized version of gkm-SVM that supports training on large-scale datasets (up to 100,000 sequences). Unlike the other tools we tested, LS-GKM uses a support vector machine instead of neural networks to detect enhancers. Additionally, this tool can be trained on a user-provided dataset, unlike the other three tools, which are distributed with pre-trained models. To train it, we randomly selected 50,000 positive sequences from the CATlas training dataset. For negative sequences, we randomly selected 10,000 sequences from each of the shuffled, LR, LNR, LGR, and LGNR training datasets, resulting in a total of 50,000 negative sequences. These 100,000 sequences were then used to train LS-GKM using its default parameters. We subsequently evaluated its performance on our testing dataset and compared the results to those of our human network. This comparison was fair, as both *EnhancerDetector* and LS-GKM were trained on the same dataset and evaluated on the same unseen testing sequences. The LS-GKM model trained on CATlas sequences was then evaluated on the 400-bp-long human and mouse ENCODE enhancers.

### 1.17 Reporter Gene Analysis

Putative enhancer sequences were synthesized as gBlocks (IDT, Coralville, IA, USA) and cloned into the plasmid pattBnucGFPs (details available upon request). Transgenic flies were generated by insertion into the third chromosome attP2 site (Rainbow Transgenics, Camarillo, CA, USA). For embryo analysis, homozygous transgenic embryos were collected, fixed, and stained using standard protocols. Larval expression was assessed by direct visualization of GFP following larval dissection.

## 2 Results

### 2.1 Main Contributions

This study presents several key contributions that advance the state of enhancer identification:

- We introduce *EnhancerDetector*, a deep learning framework for accurate and interpretable enhancer prediction across species. The code is freely available for academic use through github.com/BioinformaticsToolsmith/EnhancerDetector and doi.org/10.5281/zenodo.15531293.
- The model generalizes well to enhancers identified using diverse experimental assays, including CAGE, snATAC-seq, and DNase-seq.
- It is easily fine-tuned to new species with limited data, achieving high performance with as few as 20,000 enhancer sequences.
- Experimental validation in transgenic flies confirms the predictive accuracy of *EnhancerDetector*, with five of six tested enhancers activating reporter expression.
- We use class activation maps, an explainable AI technique that identifies the regions of an input sequence most influential to a deep network’s classification decision, together with *in silico* perturbation experiments to interpret the sequence features learned by *EnhancerDetector*. Our results show that reversing model-identified core regions disrupts enhancer predictions, revealing direction-dependent internal sequence grammar, and (ii) enhancer function emerges from an interplay between essential sequence motifs and their genomic context.
- *EnhancerDetector* consistently outperforms existing tools on both human and mouse datasets.
- Our tool identifies enhancer regions based on regulatory patterns that extend beyond simple sequence similarity.

### 2.2 *EnhancerDetector* Can Discover Human Enhancers

To assess our model’s ability to recognize human enhancers, we trained a convolutional neural network on the CATlas human tissue dataset, which includes 400-bp sequences identified as putative enhancers using snATAC-seq across 222 cell types. We evaluated several network architectures by varying the number of convolutional filters, filter sizes, and dense layer neurons. The best-performing model consisted of 64 filters in the first convolutional layer (doubled in subsequent layers) with a filter size of 3 and 20 dense neurons. As shown in Table 1, our network achieves a validation F1 score of 72%, indicating a good balance between precision and recall. The validation recall score is 66%, demonstrating that our model is able to identify about two-thirds of human enhancers. The validation precision score is 75%, meaning that for every 100 positive predictions, 75 are true positives. Our human network has high specificity—89%—indicating that it can accurately identify non-enhancers 89 out of 100 times.

**Table 1:** Training and validation performance of the human-trained convolutional network for enhancer discovery. Performance is reported on the full enhancer dataset and five control datasets: shuffled sequences, length-matched with repeats (LR), length-matched without repeats (LNR), length- and GC-matched with repeats (LGR), and length- and GC-matched without repeats (LGNR).

| Dataset | Training |  |  |  |  | Validation |  |  |  |  |
| --- | --- | --- | --- | --- | --- | --- | --- | --- | --- | --- |
|  | Accuracy (%) | Recall (%) | Specificity (%) | Precision (%) | F1 Score (%) | Accuracy (%) | Recall (%) | Specificity (%) | Precision (%) | F1 Score (%) |
| Overall | 82.83 | 69.62 | 89.44 | 76.72 | 74.18 | 81.53 | 66.34 | 89.12 | 75.30 | 72.04 |
| Shuffled | 89.24 | 69.62 | 99.05 | 97.34 | 85.91 | 88.12 | 66.38 | 98.99 | 97.05 | 84.07 |
| LR | 81.92 | 69.59 | 88.09 | 74.50 | 72.76 | 80.59 | 66.34 | 87.72 | 72.98 | 70.59 |
| LNR | 82.23 | 69.60 | 88.55 | 75.25 | 73.24 | 80.99 | 66.41 | 88.28 | 73.91 | 71.20 |
| LGR | 80.64 | 69.63 | 86.15 | 71.53 | 70.86 | 79.29 | 66.40 | 85.74 | 69.96 | 68.70 |
| LGNR | 80.09 | 69.60 | 85.33 | 70.35 | 70.07 | 78.71 | 66.34 | 84.89 | 68.70 | 67.87 |

To test the impact of control sequence characteristics, we evaluated the model against five types of negative control datasets that varied in GC content and sequence repetitiveness, along with fully shuffled enhancer sequences. When sequences were shuffled, our network achieved a high precision and specificity of 99% and 97%, respectively. These high scores show that our network can reliably distinguish actual enhancers from random sequences with the same k-mer compositions. Additionally, we evaluated the model using negative control datasets designed to control for sequence length, GC content, and the presence or absence of repetitive elements: (i) LR (length- and repeat-matched), (ii) LNR (length-matched without repeats), (iii) LGR (length- and GC-matched with repeats), and (iv) LGNR (length- and GC-matched without repeats). We observed that the network performs best when using control sequences that do not match GC content, achieving an F1 score of 71% for both the LR and LNR datasets, with LNR having a slight advantage in precision and specificity scores of 74% and 88%, respectively, compared to LR. Conversely, the model struggles more when non-enhancer sequences match GC content but do not include repeated elements, resulting in an F1 score of 68% and the lowest precision and specificity scores of 69% and 85%. Collectively, these results show *EnhancerDetector* effectively learns sequence patterns characteristic of human enhancers and can predict them with high precision and specificity.

### 2.3 *EnhancerDetector* Works on Another Vertebrate Species

To test our network’s ability to discover enhancers in other species, we assembled a dataset of putative enhancers from the mouse (*Mus musculus*) genome. This dataset allows us to evaluate whether *EnhancerDetector* can generalize to another mammalian species.

The Mouse ENCODE Project identifies putative enhancers based on DNase I hypersensitive sites. We obtained the mm10 ENCODE dataset from the UCSC Genome Table Browser. As with the human and fly enhancer datasets, we applied the same preprocessing procedure to the ENCODE dataset by removing promoters, exons, and insulators (regions containing CTCF binding sites). Promoters and exons were obtained from RefSeq and GENCODE (VM25) via the UCSC Genome Table Browser. After preprocessing, we selected potential enhancers shorter than 400 bp. Short regions were expanded bidirectionally to reach 400 bp in length. The final dataset contained 218,207 potential mouse enhancers. We then created five control datasets (shuffled, LR, LNR, LGR, and LGNR) using the same approach as for the human and fly controls. As shown in Table 2, the human-trained network achieved accuracy of 79%, recall of 66%, specificity of 85%, precision of 69%, and an F1 score of 68%. Using our base human network without any adjustments, we successfully identified two thirds of enhancers with approximately 70% precision. The model correctly excluded about 89% of non-enhancer regions. These results demonstrate that our network effectively generalizes enhancer patterns not only across different enhancer-detection biotechnologies but also across different vertebrate species.

**Table 2:** Evaluation of the human-trained network (without and with fine-tuning) on the mouse dataset. In the top half of the table, training performance is omitted because the model was not trained on mouse enhancers—it was only evaluated on mouse data.

| Dataset | Training |  |  |  |  | Validation |  |  |  |  |
| --- | --- | --- | --- | --- | --- | --- | --- | --- | --- | --- |
|  | Accuracy (%) | Recall (%) | Specificity (%) | Precision (%) | F1 Score (%) | Accuracy (%) | Recall (%) | Specificity (%) | Precision (%) | F1 Score (%) |
| *Human network with no fine-tuning* |  |  |  |  |  |  |  |  |  |  |
| Overall | — | — | — | — | — | 79.07 | 66.42 | 85.40 | 69.46 | 68.37 |
| Shuffled | — | — | — | — | — | 86.51 | 66.82 | 96.36 | 90.17 | 80.72 |
| LR | — | — | — | — | — | 77.46 | 66.79 | 82.80 | 66.01 | 66.23 |
| LNR | — | — | — | — | — | 78.15 | 66.89 | 83.77 | 67.33 | 67.14 |
| LGR | — | — | — | — | — | 77.07 | 66.95 | 82.13 | 65.19 | 65.74 |
| LGNR | — | — | — | — | — | 77.21 | 67.00 | 82.31 | 65.45 | 65.92 |
| *Human network with fine-tuning* |  |  |  |  |  |  |  |  |  |  |
| Overall | 84.39 | 75.05 | 89.06 | 77.42 | 76.45 | 83.02 | 71.72 | 88.67 | 75.99 | 74.35 |
| Shuffled | 88.24 | 75.09 | 94.82 | 87.88 | 83.13 | 86.92 | 71.79 | 94.49 | 86.69 | 81.05 |
| LR | 84.27 | 75.09 | 88.85 | 77.11 | 76.40 | 82.90 | 71.82 | 88.44 | 75.65 | 74.30 |
| LNR | 84.27 | 75.05 | 88.88 | 77.15 | 76.42 | 83.05 | 71.81 | 88.68 | 76.02 | 74.54 |
| LGR | 83.04 | 75.06 | 87.02 | 74.31 | 74.54 | 81.69 | 71.98 | 86.55 | 72.78 | 72.49 |
| LGNR | 82.16 | 75.12 | 85.68 | 72.40 | 73.25 | 80.82 | 71.79 | 85.34 | 71.00 | 71.24 |

To assess whether the predicted true positive enhancer sequences are conserved, we aligned them to the human genome using BLAST (Supplementary File 1). We found that only 6% of these sequences showed conservation (having at least 70% sequence identity and 70% alignment coverage). This low conservation rate demonstrates that our model identifies enhancer regions based on regulatory patterns that extend beyond simple sequence similarity. We next asked whether adapting the human-trained network via fine-tuning could improve performance on mouse enhancers.

### 2.4 Fine-Tuning Improves Our Mouse Enhancer Discovery

To further enhance our human network using the mouse dataset, we applied a fine-tuning technique with the potential mouse enhancers. Fine-tuning is a technique where a model that was already trained for one task is slightly adjusted to perform well on a new, related task, allowing it to keep what it has already learned while making small updates to adapt to the new data. This approach saves time and often leads to better performance, especially when the new task has less training data. We divided the enhancer and control datasets into training, validation, and testing sets using a 70%, 20%, and 10% split, respectively. To test this, we applied a fine-tuning strategy in which certain layers were frozen—their parameters remain fixed during training—while others were retrained on the mouse dataset. This technique allowed us to train a new network in which only the classifier was updated, ensuring that we retained features learned from the human enhancers while adapting to the mouse enhancers. As shown in Table 2, fine-tuning led to performance improvements across the board, compared to using the human-trained network without any adaptation. The fine-tuned network achieved an overall validation accuracy of 83%, compared to 79% for the original human network. It also showed higher recall (71% vs. 66%) and specificity (89% vs. 85%), along with improved precision and F1 scores (76% vs. 69% and 74% vs. 68%). These results highlight the effectiveness of fine-tuning in improving enhancer detection in mice, with notable gains in accuracy, recall, specificity, precision, and F1 score.

However, it is important to emphasize that even without fine-tuning, the human network remains a robust and reliable baseline. While fine-tuning provides substantial benefits when additional data is available, the human network alone offers a strong foundation for enhancer detection, delivering valuable results even in data-limited scenarios. Next, we asked: How much data are needed for effective fine-tuning?

### 2.5 The Human Network Can Be Successfully Fine-Tuned with Just 20K Mouse Enhancers

With the rise of large-scale genomic initiatives such as the 10,000 Vertebrate Genomes Project [56] and the Earth BioGenome Project [20], there is a growing demand for computational methods capable of predicting enhancers in newly sequenced genomes. Experimental approaches to enhancer identification are costly, labor-intensive, and time-consuming, making efficient *in silico* alternatives increasingly valuable. We have shown that our human-trained network generalizes reasonably well to the mouse genome without fine-tuning. However, performance improves remarkably with fine-tuning on mouse enhancer data. Additionally, certain species—whether new or well-studied—may lack a sufficient number of confirmed enhancers for effective fine-tuning. This raises a practical question: How many enhancers are needed to effectively fine-tune the human network for use in a new species? To address this question, we tested multiple datasets with varying numbers of potential enhancers required for effective fine-tuning of the human network. We used potential enhancers from the mouse training dataset, which comprised 152,744 potential enhancers, and randomly selected subsets ranging from 500 to 150,000 enhancers for training. These enhancers were sampled from the ENCODE mouse dataset and were not stratified by cell type. For each subset of enhancers, we also randomly selected twice as many control sequences from the training dataset. These subsets of enhancers and control sequences allowed us to evaluate the impact of varying numbers of potential enhancers on the fine-tuning procedure.

F1 scores on the validation set for each model are shown in Figure 4. The baseline (using the human network without fine-tuning) achieved an F1 score of 68%. Models fine-tuned on just 500 or 1,000 mouse enhancers performed comparably to or slightly worse than the baseline (66% and 68%). Marked improvements were observed when the training set included 2,000 to 20,000 enhancers, reaching up to a 73% F1 score. These gains were achieved using only about 20% of the available mouse enhancer data. While increasing the dataset size beyond 20,000 led to marginal additional gains (e.g., up to 74%), the improvements became less pronounced, suggesting that, given pooled enhancer annotations without explicit cell-type stratification, approximately 20,000 enhancers provide a strong foundation for fine-tuning the human network to a new genome. These results demonstrate the potential of our fine-tuning approach for broader cross-species enhancer prediction. We next evaluated whether *EnhancerDetector* can also learn enhancer patterns in a more distant species, *Drosophila melanogaster*.

**Figure 4:**
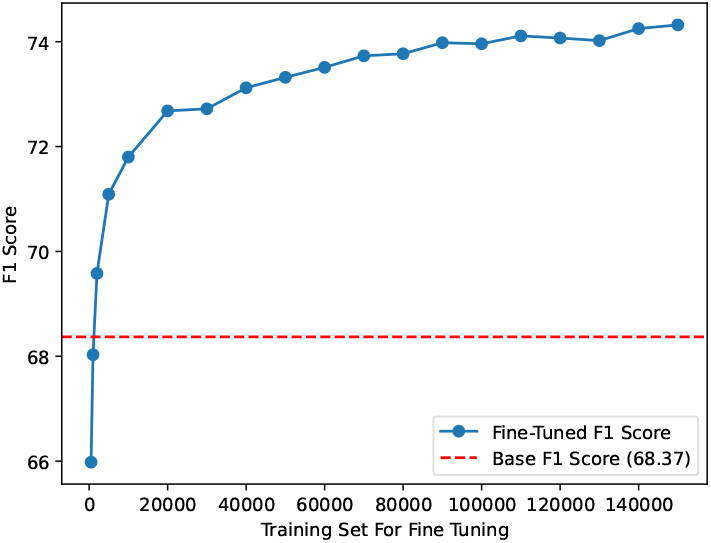
Effect of training set size on fine-tuning performance. This plot shows the F1 score achieved by the human-trained network fine-tuned on mouse enhancer data using training subsets of increasing sizes. The x-axis indicates the number of mouse enhancer sequences used for fine-tuning, while the y-axis reports the corresponding F1 score on the validation set. Increasing the number of fine-tuning samples improves enhancer detection performance, with performance gains plateauing at larger dataset sizes.

### 2.6 *EnhancerDetector* Can Learn Patterns in Fly Enhancers

To evaluate *EnhancerDetector* ‘s ability to identify enhancers in *Drosophila melanogaster*, we built multiple convolutional networks with varying numbers of filters, filter sizes, and dense layer neurons. We selected the two models that gave the best results (Table 3). These networks were trained using a combination of fly enhancer datasets: distal DNase I hypersensitive sites, distal scATAC-seq peaks, and cis-regulatory modules from REDFly. As with the human and mouse experiments, we used five negative datasets. Overall, both models had 67–68% validation F1 scores. Their validation recall scores ranged 63–68%, and specificity scores ranged 89–91%. Their precision scores were 67–70%. The scores on the training data were slightly higher than those obtained on the validation dataset but still comparable, demonstrating the lack of overfitting. The two networks performed well on the shuffled dataset. Both models showed strong specificity and precision when the control sequences did not contain repeats. Controlling for GC content made the classification task slightly more difficult: the F1 scores of the two networks on the LNR dataset (control for length only, excluding repeats) were both 68%, whereas the scores on the LGNR (length and GC-controlled, no repeats) dataset were both 67%. The same trend—that controlling for GC content makes the task slightly more difficult—was observed on the datasets that include repeats. Compared to the performance of the human network, the trend related to GC content is consistent, but the trend related to the inclusion or exclusion of repeats is not. These results reaffirm previous findings [57] that enhancer sequences exhibit a discernible pattern and are characterized by a specific GC content.

**Table 3:** Training and validation performance of different neural networks on the fly enhancers. The models include two fly-trained convolutional networks, a fine-tuned human convolutional network on fly data, an ensemble combining ab initio and fine-tuned models for discovering fly enhancers, and a recurrent model specifically designed for fly enhancer sequences. Performance was also evaluated on five control datasets: four sets consisting of genomic sequences sampled to match the enhancer datasets in length and/or GC content, with or without repetitive elements (LR: Length Repeat; LNR: Length No Repeat; LGR: Length GC Repeat; LGNR: Length GC No Repeat), and a fifth set consisting of shuffled enhancer sequences.

| Dataset | Training |  |  |  |  | Validation |  |  |  |  |
| --- | --- | --- | --- | --- | --- | --- | --- | --- | --- | --- |
|  | Accuracy (%) | Recall (%) | Specificity (%) | Precision (%) | F1 Score (%) | Accuracy (%) | Recall (%) | Specificity (%) | Precision (%) | F1 Score (%) |
| *Fly from Scratch (32 filters of size 3 and 20 dense units)* |  |  |  |  |  |  |  |  |  |  |
| Overall | 85.96 | 66.72 | 92.38 | 74.47 | 71.67 | 83.82 | 68.45 | 88.94 | 67.35 | 67.69 |
| Shuffled | 90.63 | 66.75 | 98.59 | 94.05 | 82.73 | 90.88 | 68.49 | 98.35 | 93.25 | 83.19 |
| LR | 84.25 | 66.69 | 90.10 | 69.18 | 68.29 | 80.89 | 68.51 | 85.02 | 60.38 | 62.83 |
| LNR | 85.85 | 66.73 | 92.22 | 74.09 | 71.42 | 84.04 | 68.38 | 89.27 | 67.98 | 68.10 |
| LGR | 83.59 | 66.70 | 89.23 | 67.35 | 67.09 | 79.97 | 68.46 | 83.81 | 58.49 | 61.44 |
| LGNR | 85.47 | 66.70 | 91.73 | 72.90 | 70.67 | 83.28 | 68.47 | 88.23 | 65.96 | 66.74 |
| *Fly from Scratch (40 filters of size 3 and 20 dense units)* |  |  |  |  |  |  |  |  |  |  |
| Overall | 84.92 | 59.21 | 93.49 | 75.21 | 68.96 | 83.94 | 62.92 | 90.94 | 69.84 | 67.34 |
| Shuffled | 89.44 | 59.22 | 99.52 | 97.61 | 80.21 | 90.26 | 62.91 | 99.37 | 97.09 | 82.16 |
| LR | 83.25 | 59.20 | 91.26 | 69.31 | 65.52 | 81.27 | 62.96 | 87.38 | 62.42 | 62.57 |
| LNR | 84.80 | 59.23 | 93.32 | 74.73 | 68.68 | 84.21 | 62.91 | 91.31 | 70.70 | 67.86 |
| LGR | 82.68 | 59.21 | 90.50 | 67.52 | 64.45 | 80.48 | 62.99 | 86.32 | 60.53 | 61.30 |
| LGNR | 84.48 | 59.28 | 92.89 | 73.53 | 68.02 | 83.54 | 62.96 | 90.40 | 68.60 | 66.59 |
| *Human Fine-tuned on Fly* |  |  |  |  |  |  |  |  |  |  |
| Overall | 87.68 | 73.24 | 92.49 | 76.47 | 75.13 | 83.27 | 69.36 | 87.90 | 65.64 | 66.60 |
| Shuffled | 92.14 | 73.24 | 98.44 | 93.99 | 85.85 | 90.92 | 69.34 | 98.11 | 92.45 | 83.15 |
| LR | 86.04 | 73.25 | 90.31 | 71.58 | 72.10 | 80.14 | 69.40 | 83.73 | 58.70 | 61.83 |
| LNR | 87.61 | 73.26 | 92.39 | 76.24 | 75.18 | 83.49 | 69.47 | 88.16 | 66.18 | 67.20 |
| LGR | 85.39 | 73.25 | 89.43 | 69.80 | 70.87 | 79.14 | 69.40 | 82.39 | 56.77 | 60.38 |
| LGNR | 87.25 | 73.29 | 91.90 | 75.10 | 74.45 | 82.64 | 69.42 | 87.05 | 64.11 | 65.75 |
| *Ensemble of ab initio and Fine-tuned on Fly* |  |  |  |  |  |  |  |  |  |  |
| Overall | 85.39 | 52.14 | 96.47 | 83.13 | 64.09 | 83.85 | 52.97 | 94.15 | 75.10 | 62.13 |
| Shuffled | 87.95 | 52.14 | 99.89 | 99.35 | 68.39 | 88.12 | 52.97 | 99.84 | 99.08 | 69.04 |
| LR | 84.33 | 52.14 | 95.06 | 77.88 | 62.46 | 81.95 | 52.97 | 91.62 | 67.80 | 59.48 |
| LNR | 85.43 | 52.14 | 96.52 | 83.32 | 64.14 | 84.11 | 52.97 | 94.48 | 76.20 | 62.50 |
| LGR | 84.00 | 52.14 | 94.62 | 76.36 | 61.97 | 81.40 | 52.97 | 90.87 | 65.92 | 58.74 |
| LGNR | 85.24 | 52.14 | 96.27 | 82.33 | 63.85 | 83.68 | 52.97 | 93.92 | 74.39 | 61.88 |
| *Recurrent Model for Fly Enhancers* |  |  |  |  |  |  |  |  |  |  |
| Overall | 84.15 | 61.92 | 91.56 | 70.96 | 67.64 | 83.62 | 67.72 | 88.92 | 67.08 | 67.26 |
| Shuffled | 89.82 | 61.90 | 99.13 | 95.95 | 81.04 | 91.14 | 67.73 | 98.94 | 95.53 | 84.00 |
| LR | 82.27 | 61.90 | 89.04 | 65.33 | 64.11 | 80.71 | 67.70 | 85.04 | 60.14 | 62.41 |
| LNR | 83.76 | 61.89 | 91.05 | 69.74 | 66.87 | 83.71 | 67.66 | 89.05 | 67.33 | 67.38 |
| LGR | 81.52 | 61.93 | 88.04 | 63.32 | 62.81 | 79.79 | 67.77 | 83.79 | 58.23 | 61.06 |
| LGNR | 83.37 | 61.93 | 90.51 | 68.51 | 66.12 | 82.82 | 67.72 | 87.86 | 65.02 | 65.87 |

### 2.7 Fine-Tuning the Human Network Achieved Comparable Performance to Fly Networks Trained Ab Initio

Having established strong *ab initio* performance on fly enhancers, we next tested whether a human-trained model can be adapted to fly via fine-tuning. Because the human training dataset was substantially larger than the fly dataset (8.9 million vs. 6.4 million samples), we hypothesized that the human-trained model had captured generalizable sequence features. Similar to our approach with the mouse dataset, we froze all layers during training except for the dense layers, which are responsible for making classifications rather than extracting features. This approach allowed the model to retain previously learned representations while adapting to fly-specific enhancer sequence patterns. As shown in Table 3, the fine-tuned network performed comparably to the two models trained from scratch on the fly dataset. The overall validation accuracy was 83%, compared to 84% for both fly-specific models. The fine-tuned network achieved a recall of 69%, slightly outperforming the two *ab initio* models (68% and 63%), while maintaining strong specificity (88% compared to 89% and 91%). Precision (66%) and F1 score (67%) were also closely aligned with those of the fly-trained models. These results demonstrate that fine-tuning a human-trained model can be as effective as training a model *ab initio*, offering a practical and efficient strategy for adapting *EnhancerDetector* to new species without the need for extensive retraining or large datasets.

### 2.8 A Network Ensemble Improves Prediction Confidence

After training three fly models—two trained from scratch on the fly dataset and one fine-tuned from the human-trained network—we observed that their individual performances were comparable, yet slightly different. To improve prediction confidence, we combined these models into an ensemble. In this approach, each model independently evaluates a sequence, and the sequence is classified as an enhancer only if all three models agree. This voting-based strategy prioritizes precision and specificity over recall by requiring unanimous consensus across models when it comes to positive predictions. As shown in Table 3, the ensemble achieved a specificity score of 94% and a precision score of 75%, outperforming the individual models (specificity scores of 89%, 91%, and 88%; precision scores of 67%, 70%, and 66%). Although recall dropped to 53%, indicating that the ensemble identified fewer true enhancers, it substantially reduced the number of false positives. These results demonstrate that the ensemble method provides a valuable option when higher confidence in predicted enhancers is needed, such as for experimental validation using transgenic animals.

### 2.9 *EnhancerDetector* Can Determine Enhancer Locations Within Longer Sequence Fragments

We selected 18 long sequences (median length 4 kb, range 3.2–7.1 kb) from the REDfly database. Each of these sequences has demonstrated regulatory activity and thus contains one or more individual enhancers. Our goal was to determine the locations of the enhancers within these long sequences. We cleaned the sequences by removing promoters, insulators, and exons (see Methods). We then split each long sequence into segments ranging from 460 to 500 base pairs and fed each segment into our ensemble of networks. Based on the confidence scores generated by the networks, we determined the locations of the highest-scoring segment(s) within each long sequence. We also shuffled the long sequences and ran them through the ensemble to ensure that our network could distinguish between real enhancer sequences and shuffled sequences. To test whether the network ensemble accurately predicted discrete enhancer locations within the longer sequences, we selected six predicted enhancers for *in vivo* testing using reporter gene analysis in transgenic *Drosophila*. Results are shown in Figure 5. Three of the putative enhancers (HPC1–HPC3) were drawn from the same initial long sequence, while three others were taken from distinct long sequences (HPC4, HPC7, and HPC8). HPC1 drives weak reporter gene expression in the embryonic midgut and dorsal ectoderm (Figure 5 A, B), consistent with the activity of its enclosing longer sequence, enhancer ush -12475/-6925 (REDflyID: RFRC:0000003436.004; Muratoglu et al. [58]). Interestingly, as reported in the original study [58], this expression appears to be ectopic relative to the endogenous expression of the assigned target gene, *ush*. HPC2 drives reporter gene expression in the midline of the embryonic ventral nervous system (Figure 5 C). This expression is not observed with the original longer enhancer. However, it is consistent with the known expression pattern of the *ush* gene, although a precise one-to-one correspondence between expressing cells remains to be determined [59]. Whether the discrepancy between the two reporter constructs results from a mis-characterization of the original ush_- 12475/-6925 reporter, is the result of a latent activity of the HPC2 fragment that is masked either *in vivo* or in the removed-from-context longer fragment, or is due to some other reason similarly remains to be explored. HPC3 failed to show reporter activity (data not shown). HPC4 exhibited strong reporter activity in the embryonic salivary gland (Figure 5 D). However, this is not a tissue where activity was observed with the original longer sequence (GMR26G07; REDflyID: RFRC:0000024999.002; Jenett et al. [14]).

**Figure 5:**
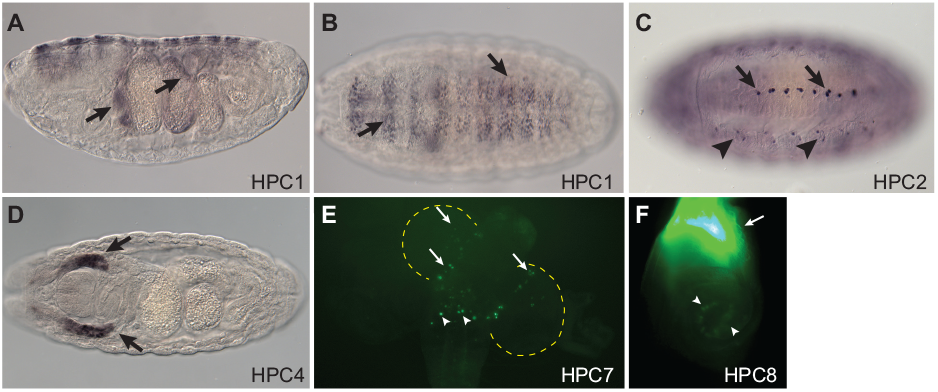
*In vivo* validation of predicted enhancers. *Drosophila* enhancers of 460–500 bp in length were predicted from longer sequences (about 4,000 bp) with known regulatory activity and tested *in vivo* in a transgenic reporter gene assay. (A) HPC1 drives expression in the embryonic midgut (arrows) as well as (B) the dorsal ectoderm (arrows), consistent with described expression for the enclosing ush -12475/-6925 enhancer [58]. (C) HPC2 drives reporter gene expression in the ventral midline (arrows), consistent with known expression of the *ush* gene but not the ush -12475/-6925 enhancer. Arrowheads indicate expression driven by the plasmid vector, not the HPC2 enhancer. (D) HPC4 shows reporter gene expression in the embryonic salivary glands (arrows). Neither the larger enclosing enhancer (GMR26G07) nor its assigned target gene (*chn*) are reported to be expressed in this tissue. (E) HPC7 drives reporter expression in the larval brain (arrows; brain is outlined with yellow dotted lines) and ventral nerve cord (arrowheads). This resembles the described expression of the enclosing GMR52B12 enhancer. (F) HPC8 shows reporter gene expression in imaginal discs, including the leg discs (pictured). Very strong expression is observed in the presumptive proximal leg tissue (arrow), with weaker expression more distally (arrowheads), consistent with described expression of the *hth* target gene.

Moreover, ectopic expression in this tissue is not uncommon due to sequences in the reporter gene vector [60]. Thus, it is questionable whether this sequence should be considered a *bona fide* enhancer. HPC7 did not show significant embryonic activity but did drive limited gene expression in the larval brain and ventral nerve cord (Figure 5 E). This is consistent with the described activity of the longer enclosing sequence (GMR52B12; REDflyID: RFRC:0000027000; Jenett et al. [14]), although, again, an exact one-to-one correspondence has not been determined. HPC8 also lacked appreciable embryonic activity but was active in imaginal discs (Figure 5 F), consistent with part of the described activity of its longer enclosing sequence (GMR46A08; REDflyID: RFRC:0000030377; Jenett et al. [14]). In total, five of the six tested putative enhancers (83%) drove reporter gene activity, and four (67%) exhibited expression patterns consistent with those of their larger enclosing sequences and/or their likely target genes. These results strongly demonstrate that *EnhancerDetector* can successfully identify transcriptional enhancers.

### 2.10 Recurrent and Convolutional Networks Have Comparable Performance

To evaluate alternative architectures for enhancer prediction, we trained a Recurrent Neural Network (RNN) from scratch on the fly dataset and compared its performance to that of our three Convolutional Neural Networks (CNNs): two trained directly on fly data and one fine-tuned from the human-trained model. The RNN architecture employed gated recurrent units and followed a structure similar to the CNNs, including an embedding layer, two recurrent blocks, and two dense layers. As shown in Table 3, the RNN achieved validation performance comparable to that of the CNNs. Overall, the validation accuracy scores for the four networks were approximately the same (84% vs. 84%, 84%, and 83%). Recall scores were also similar (68% vs. 68%, 63%, and 69%), as were specificity (89% vs. 89%, 91%, and 88%) and precision (67% vs. 67%, 70%, and 66%). Importantly, the F1 scores—the harmonic mean of recall and precision— were nearly identical across all models (67% vs. 68%, 67%, and 67%). These results demonstrate that the RNN achieves performance comparable to the CNNs, indicating that both architectures are suitable for recognizing transcriptional enhancers. However, training the RNN was time-consuming. Each epoch required approximately 40 minutes, and training the network on the fly data took around 65 epochs, resulting in a total training time of about two days. In contrast, each epoch of the CNN took about 8 minutes, and training typically required 22 epochs on the fly data—resulting in a total training time of approximately three hours. Thus, the CNNs trained markedly faster while achieving similar performance. When we attempted to train an RNN on the human dataset, each epoch took nearly four hours, rendering the training process impractically slow. For this reason, we did not pursue RNN training on the human data. In summary, our decision to prioritize CNNs over RNNs was clearly justified: they achieved comparable performance while drastically reducing the required training time.

### 2.11 *EnhancerDetector* Detects Putative Enhancers Identified by Alternative Biotechnologies

To evaluate our human network, we used an alternative set of potential enhancers identified using different methodologies. The FANTOM5 Project identifies putative enhancers based on cap analysis of gene expression, a method distinct from the snATAC-seq technology used to identify CATlas enhancers. We applied the same preprocessing pipeline to the FANTOM dataset as we did for the fly and human datasets, including the removal of promoters, exons, and insulators. Next, we extracted regions that are 400 bp or shorter; any region shorter than 400 bp was symmetrically expanded to 400 bp. This step was necessary because the human network—trained on CATlas enhancers—expects input sequences of 400 bp. After preprocessing, the dataset contained 41,553 potential enhancers. We first performed a baseline evaluation using our human network to assess its general ability to detect enhancers. To ensure a fair assessment, we then divided the FANTOM5 regions into four datasets: training, validation, testing, and non-overlapping. A region was assigned to the training, validation, or testing set if it overlapped a CATlas region in the corresponding set by at least 50%. The non-overlapping set included FANTOM regions that did not meet this overlap threshold with any CATlas training, validation, or testing region. This partition procedure resulted in 17,306, 4,896, 2,483, and 16,868 sequences in the respective datasets.

When evaluated on the full FANTOM5 dataset, the model achieved an accuracy of 73%, a recall of 50%, a specificity of 96%, and a precision of 93%. Although the recall was lower than in the CATlas-based evaluations, the high precision and specificity suggest that the model is stringent in its predictions, favoring precision over sensitivity and thereby reducing false positives. On the training, validation, and testing subsets of the FANTOM5 data, the network correctly identified 66–67% of the putative enhancers. Although the FANTOM5 testing subset contains some regions overlapping with CATlas, it is important to emphasize that the network had no access to this data during training or validation; hence, results on this set represent a true blind test. Detecting 67% of the putative enhancers in such a blind evaluation underscores our model’s ability to generalize enhancer patterns across datasets.

On the non-overlapping subset—which contains sequences with no or minimal (*<*50%) overlap with CATlas—the network classified only 28% of the sequences as enhancers. While a 28% detection rate for the non-overlapping set may initially appear low, it is important to note that approximately 25–30% of sequences in the FANTOM5 dataset are not true enhancers. As reported by the FANTOM5 authors, “67.4–73.9% of the CAGE-defined enhancers showed significant reporter activity.” In another *in vivo* validation using zebrafish embryos, they observed “tissue-specific enhancer activity with 3 of 5 fragments, which corresponded to the human enhancer tissue expression.” Thus, the 28% recall rate on the non-overlapping set doesn’t necessarily reflect poor model performance, but rather suggests that this subset predominantly comprises the false positives within the FANTOM5 dataset, a hypothesis strongly supported by their minimal or absent overlap with the high-confidence single-cell data from CATlas.

These results demonstrate that our network effectively identifies a substantial portion of enhancers detected via both CAGE and snATAC-seq, highlighting its generalizability across datasets produced using different biotechnologies.

### 2.12 CAM-Based Knockout Experiments Reveal Biologically Significant Enhancer Regions

Class Activation Maps (CAMs) are visual tools that highlight the regions of an input sequence that are most influential in a neural network’s classification decision. They help interpret which features the model has learned to associate with a given class. This technique outputs a score for each nucleotide in the sequence, typically ranging from 0 (least important) to 1 (most important). These scores can be visualized as a heat map: nucleotides with scores closer to 1 appear reddish, indicating high importance, while those closer to 0 appear bluish, indicating low importance. An example CAM is shown in Figure 6. In this figure, the dark red region spanning positions 110 to 140 highlights a portion of the sequence that strongly influences the network’s classification. The green region spanning positions 260 to 290 highlights another portion of the sequence that moderately influences the network’s classification. In contrast, the blue regions indicate areas with little to no relevance to the network’s decision-making process.

**Figure 6:**
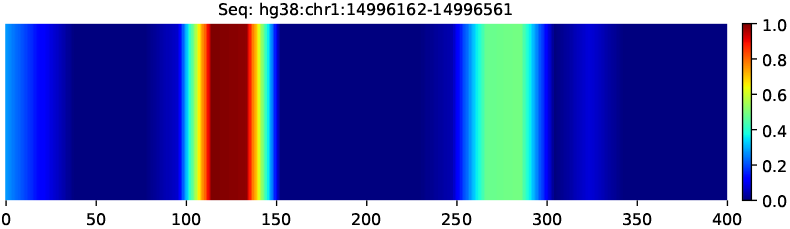
Class Activation Map (CAM) visualization for a true enhancer—a sequence labeled as an enhancer and correctly classified as such by the neural network. CAM is a technique that assigns importance scores to each nucleotide in the sequence based on how much it contributed to the model’s classification. These scores range from 0 (least important) to 1 (most important) and are visualized as a heat map: nucleotides with scores closer to 1 appear reddish, while those closer to 0 appear bluish. In this example, the dark red region spanning positions 110 to 140 indicates a strong enhancer signal and the green region spanning positions 260 to 290 indicates a moderate enhancer signal, suggesting that these portions of the sequence played a key role in the network’s decision. The blue regions were less or not influential at all.

To evaluate the biological significance of the regions highlighted by CAMs, we conducted a knockout experiment. The key idea was to disrupt (“knock out”) the most important region of an enhancer sequence— identified using CAM—and assess whether the sequence was still classified as an enhancer by the network. We began with 25,000 true enhancer sequences and computed their CAM scores to identify regions of high importance. We defined important regions using a thresholding system with values of 0.2, 0.3, 0.4, and 0.5. These thresholds represent the minimum average CAM score required for a region to be considered important. Based on this criterion, we selected sequences containing only one important region between 25 and 100 bp in length. This constraint was critical: destroying long important regions would disrupt a large portion of the sequence and likely lead to a negative classification, while regions shorter than 25 bp would be too small to meaningfully impact classification. For each selected sequence, we shuffled the nucleotides within the important region, effectively removing its recognizable pattern. As a control, we shuffled a region of the same length elsewhere in the sequence—ensuring it did not overlap with the important region. The classifier was then applied to modified sequences to observe the effect of the knockouts.

Finally, we assessed the significance of the results using a binomial test, which calculates p-values based on the number of sequences tested. This statistical evaluation helped confirm the observed differences in classification outcomes were not due to random chance, but rather to the disruption of biologically relevant regions identified by CAMs.

Shuffling the regions identified as important by CAMs significantly reduced the number of sequences classified as enhancers across all thresholds. At a threshold of 0.2, 1,409 sequences (94% of true enhancers with important regions averaging an intensity of 0.2) were reclassified as non-enhancers. This effect persisted across higher thresholds: 2,251 sequences (92%) at 0.3, 3,070 (90%) at 0.4, and 4,345 (87%) at 0.5. In total, disrupting CAM-identified regions led to loss of enhancer classification in 87–94% of cases. As a control, we shuffled a region outside the CAM-highlighted area within the same sequences. This had a smaller but still substantial effect: at thresholds of 0.2, 0.3, 0.4, and 0.5, the number of sequences reclassified as non-enhancers was 1,281 (85%), 1,928 (79%), 2,443 (72%), and 3,371 (68%). Next, we computed p-values comparing the proportion of negative classifications resulting from shuffling important versus non-important regions. The p-values for thresholds 0.2 through 0.5 were 6.87 *×*10^*−*26^, 8.94 *×*10^*−*71^, 1.34 *×*10^*−*151^, and 1.08 *×* 10^*−*225^—all highly significant. We also evaluated two alternative knockout strategies: reverse sequencing (e.g., AGCT*→* TCGA) and transversion mutation, where each nucleotide (A, C, G, T) is randomly replaced with either of the two remaining non-complementary bases (e.g., A becomes G or C, determined by chance). Both approaches yielded reclassification rates comparable to the shuffling approach. At a threshold of 0.2, the reclassification rates to non-enhancer were 92% for the mutation approach and 95% for reverse sequencing vs. 94% for the shuffled knockout. At 0.3, the rates were 89% and 92% vs. 92%. For 0.4, we observed 86% and 91% vs. 90%. Finally, at 0.5, the rates were 84% and 88% vs. 87%. All of these percentages were statically significant.

These results demonstrate that the regions identified by CAMs are not only predictive but also biologically meaningful. Disrupting the important regions has a greater impact on enhancer classification than disturbing other regions. Notably, the experimental reversal of key enhancer regions strongly suggests a directional nature, independently confirming recent observations [61].

### 2.13 Insertion Experiment Further Supports the Biological Relevance of CAM-Identified Regions

To further demonstrate the importance of the CAM-identified regions, we conducted an insertion experiment. The goal was to test whether inserting a high-scoring region from a true enhancer into a non-enhancer sequence could cause the network to reclassify it as an enhancer. We began with 25,000 true enhancer sequences and applied the same selection criteria as in the knockout experiment: each sequence had to contain exactly one important region identified by CAM, with a length between 25 and 100 bp. For each selected enhancer, we paired it with a true negative sequence—one that was labeled and correctly classified as a non-enhancer. We then inserted the important region from the enhancer into the corresponding negative sequence at the same relative position. After this modification, the new sequence was passed through the network for reclassification. As a control, we performed a parallel experiment in which a randomly selected region of the same size–but from outside the CAM-highlighted area–was inserted into the negative sequence in the same location. This ensured that any observed changes in classification were specifically due to the informative region identified by CAM, rather than arbitrary sequence changes. Finally, we evaluated the significance of the observed classification changes using a binomial test, demonstrating that the effects were not due to random chance.

For CAM thresholds of 0.2, 0.3, 0.4, and 0.5, the number of sequences reclassified as enhancers was 1,132 (76%), 1,937 (79%), 2,745 (80%), and 4,030 (81%). As a control, we inserted regions outside of the CAM-identified important areas into true negatives. The number of sequences reclassified as enhancers in this case was 877 (59%), 1,485 (61%), 2,042 (60%), and 2,990 (60%) for the respective thresholds. We then calculated p-values comparing the reclassification rates resulting from inserting important versus non-important regions. The p-values for thresholds 0.2 through 0.5 were 1.70 *×* 10^*−*43^, 4.01 *×* 10^*−*85^, 3.87*×*10^*−*146^, and 2.65*×*10^*−*220^—all highly significant. Insertion of CAM-identified regions consistently increased enhancer classification rates across all thresholds, with stronger effects at higher CAM intensities. In contrast, insertion of non-important regions had moderate but notably weaker impact. These findings confirm the biological relevance of CAM-identified regions in enhancer sequences.

### 2.14 Sequence Context Surrounding Enhancer Regions Contributes to Classification

While our knockout and insertion experiments demonstrated the functional significance of CAM-identified regions, we next examined whether the surrounding sequence context also influences enhancer classification. Specifically, we asked: Is preserving only the important region sufficient for the model to recognize a sequence as an enhancer? We designed the experiment to be similar to the knockout experiment, except instead of shuffling the important region itself, we shuffled the adjacent context of same length as the important region. For example, if the important region is 100 base pairs long, we shuffle the 50 base pairs directly before and after it, while keeping the important region unchanged. This allowed us to test whether the network relies on the broader sequence context, beyond the specific important region, for enhancer classification. As with the previous experiments, we selected 25,000 true enhancer sequences, applying the same criteria for selection: each sequence contained one important region between 25 and 100 bp in length. Once the sequences were chosen, we took the length of the important region and shuffled the regions on both adjacent sides of the important region to match. After modifying the sequences, we passed them through the network again to determine whether they would still be classified as enhancers.

At a CAM threshold of 0.2, 1,277 sequences (85% of the samples) were reclassified as non-enhancers. For thresholds of 0.3 and 0.4, the number of reclassified sequences was 1,959 (80%) and 2,524 (74%). At threshold 0.5, 3,553 sequences (71%) were reclassified as non-enhancers. Similar to the knockout experiment, we applied the following three disturbance approaches: (i) k-mer shuffling, (ii) transverse mutation, and (iii) sequence reverse. The results were comparable to those of the shuffled approach. At the 0.2 threshold, reclassification rates were 82% for mutation, 86% for reverse, and 85% for shuffled. At 0.3, the rates were 76% and 81% vs. 80%. For 0.4, we observed 71% and 75% vs. 74%. Finally, at 0.5, the rates were 69% and 74% vs. 71%.

These results indicate that preserving only the most predictive region is insufficient for accurate enhancer classification. Disrupting the surrounding sequence context often leads the model to reject the sequence as an enhancer, even when the core region remains intact. This finding is consistent with biological evidence suggesting that enhancer function depends not only on core transcription factor binding motifs but also on adjacent nucleotides that influence chromatin accessibility and higher-order regulatory architecture [62].

A comparison with the knockout experiments reveals that both the core region and its context contribute to enhancer activity. However, perturbing the core region resulted in a higher rate of loss-of-function predictions, indicating it plays a more central role. Together, these findings suggest that enhancer function emerges from an interplay between essential sequence motifs and their genomic context—both of which are effectively captured by our model.

### 2.15 *EnhancerDetector* Outperforms LS-GKM and DeepSEA on the CATlas Testing Set

We evaluated *EnhancerDetector* on the held-out CATlas test set, comparing its performance with the k-mer SVM LS-GKM and the chromatin-feature predictor DeepSEA. We could not evaluate other related tools—e.g., Enhancer-FRL, Enhancer-LSTMAtt, and PDCNN—on the CATlas dataset because these tools require 200-bp-long input sequences, whereas the CATlas sequences are 400 bp long; downsampling to 200 bp would likely remove informative context and yield an unfair comparison. All CATlas evaluations are based on a test set with a 1:10 enhancer-to-control ratio. The results are shown in Figure 7.

**Figure 7:**
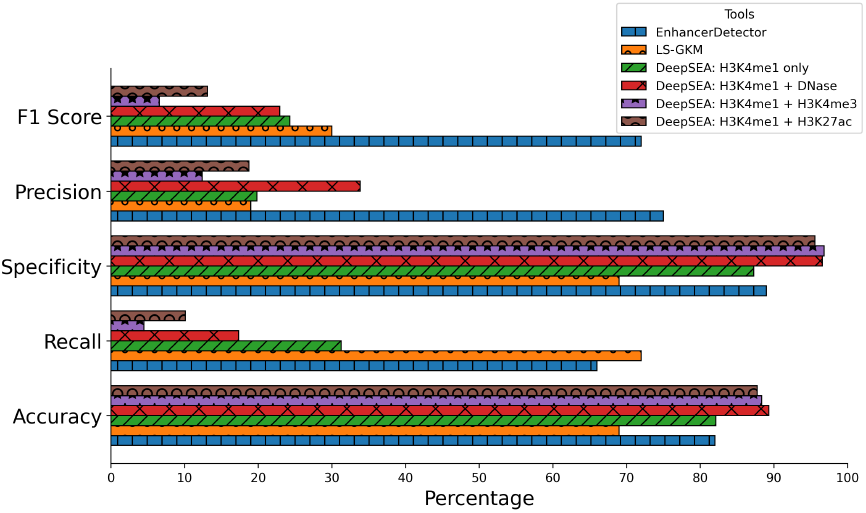
Performance of *EnhancerDetector* compared with LS-GKM and DeepSEA on the CATlas human test set. Grouped horizontal bars showed accuracy, recall, specificity, precision, and F1 score for each tool. *EnhancerDetector* achieved the highest precision and F1 score while also maintaining strong recall and specificity. LS-GKM identified more positives but at the cost of substantially reduced precision. DeepSEA, evaluated using four chromatin-mark rules, was conservative, showing high specificity but low recall and precision.

We first compared our network to LS-GKM trained on the CATlas dataset, using the same unseen test set for evaluation. Our model outperformed LS-GKM in overall accuracy (82% vs. 69%). While LS-GKM achieved higher recall (72% vs. 66%), our model showed superior specificity (89% vs. 69%), precision (75% vs. 19%), and F1 score (72% vs. 30%). These results suggest that LS-GKM tends to over-classify sequences as enhancers, whereas *EnhancerDetector* achieves a more balanced performance, making it better suited for genome-scale enhancer discovery. To further assess performance beyond single-point metrics, we generated receiver operating characteristic (ROC) and precision-recall (PRC) curves. When trained with 50,000 enhancer and 50,000 control sequences, LS-GKM achieved an AUROC of 0.775 and an AUPRC of 0.275, reflecting its higher recall but limited precision. In contrast, *EnhancerDetector* achieved an AUROC of 0.879 and a substantially higher AUPRC of 0.507, demonstrating a more favorable balance between sensitivity and precision particularly important in the context of highly imbalanced genomic data.

LS-GKM is based on support vector machines, which are known for their limited ability to scale on large data sets. For this reason, we further optimized LS-GKM; we trained additional models using progressively smaller subsets of the CATlas training data (ranging from 10k to 40k enhancers and controls). We observed a steady improvement in both AUROC and AUPRC with increasing training size: AUROC rose from 0.749 (10k) to 0.771 (40k), while AUPRC increased from 0.239 to 0.269. Despite these gains, the 50k model remained the best-performing LS-GKM configuration yet still did not surpass *EnhancerDetector*. We did not exceed 50k samples, as the LS-GKM documentation recommends a maximum of 100,000 total sequences, and we aimed to remain within that constraint.

We then compared to DeepSEA, a widely used deep learning framework designed for regulatory variant prediction [42], specifically to their Beluga model [63]. While *EnhancerDetector* can distinguish enhancers from controls using direct sequence classification, we sought to assess how it performs relative to an alternative approach that infers enhancer activity through chromatin feature prediction. DeepSEA is not a dedicated enhancer detector; rather, it predicts cell-type specific open chromatin features such as DNase I hypersensitivity and histone modifications (e.g., H3K4me1, H3K27ac, H3K4me3) from DNA sequences. In this section, we evaluate DeepSEA on the CATlas human test set to enable a controlled comparison under identical conditions. In the next section, we extend this benchmarking framework to independent ENCODE human and mouse enhancer datasets.

Since DeepSEA requires 2000-bp-long input sequences, we symmetrically extended each 400 bp region to 2,000 bp by adding flanking sequence on both sides, keeping the original enhancer region centered. To interpret DeepSEA’s predictions in the context of enhancer classification, we defined enhancers based on the presence of key chromatin marks. Specifically, we considered four scenarios: (1) H3K4me1 alone, a primary enhancer-associated mark; (2) H3K4me1 + DNase, reflecting accessible chromatin surrounding enhancer elements; (3) H3K4me1 + H3K27ac, indicative of active enhancers; and (4) H3K4me1 + H3K4me3, while traditionally associated with promoters, has shown signals in enhancers although less than H3K4me1 [64]. We defined a region as an enhancer if the predicted probability for any combination exceeded 0.5, regardless of cell type. This cell-agnostic thresholding reflects our focus on general enhancer potential rather than cell-specific activity.

We evaluated DeepSEA on the CATlas human test set. Across all chromatin mark combinations, DeepSEA achieved an overall accuracy ranging from 82% to 89% (H3K4me1 alone and H3K4me1 + DNase), with recall ranging from as low as 4% (for H3K4me1 + H3K4me3) to 31% (for H3K4me1 alone). Specificity ranged from 87% (H3K4me1 alone) to 97% (H3K4me1 + H3K4me3), indicating the model’s tendency to classify sequences as negatives. Precision and F1 score were both low overall, for precision ranging from 12% (H3K4me1 + H3K4me3) to 20% (H3K4me1 alone) and F1 score ranging from 6% (H3K4me1 + H3K4me3) to 24% (H3K4me1 alone). In contrast, *EnhancerDetector* achieved substantially higher performance on recall, precision and F1 score while maintaining high specificity, with 82% accuracy, 66% recall, 89% specificity, 75% precision, and 72% F1 score. These results show that while DeepSEA effectively captures chromatin-associated signals from DNA sequence, its indirect approach to enhancer inference may limit its sensitivity for this specific task. In contrast, *EnhancerDetector* is optimized for enhancer classification and provides the best compromise between recall and precision, as evidenced by the highest F1 score, making it well suited for direct enhancer detection.

### 2.16 *EnhancerDetector* Outperforms Multiple Enhancer-Discovery Tools on Independent ENCODE Datasets

To evaluate performance on an independent benchmark distinct from CATlas, we next compared our human-trained network to multiple tools for enhancer classification on ENCODE human and mouse datasets. We compared our human-trained network to several other binary classification tools for human enhancer prediction: Enhancer-FRL, Enhancer-LSTMAtt, PD-CNN, and LS-GKM (unlike the previous section, where LS-GKM was evaluated on CATlas, here all tools were evaluated on ENCODE-derived enhancer datasets based on DNase I hypersensitivity). We evaluated all tools on the ENCODE datasets for both human and mouse (Figure 8). All evaluations were performed on a test set with a 1:10 enhancer-to-control ratio, reflecting the natural sparsity of enhancers and imposing a strong penalty on precision and F1 score. Against LS-GKM (trained on CATlas but evaluated here on ENCODE), our model achieved higher accuracy (87% vs. 70%), specificity (90% vs. 70%), precision (35% vs. 18%), and F1 score (42% vs. 29%) on the human data, though LS-GKM retained higher recall (66% vs. 54%). On the mouse data, LS-GKM again showed higher recall (72% vs. 62%) but substantially lower precision (16% vs. 30%) and F1 (26% vs. 40%). Enhancer-FRL and Enhancer-LSTMAtt displayed a similar tradeoff: both achieved higher recall but much lower precision and specificity. For example, Enhancer-FRL reached 62% recall on the human data but only 9% precision, compared to our model’s 52% recall and 35% precision (F1: 13% vs. 42%). Enhancer-LSTMAtt likewise showed higher recall on the mouse data (77% vs. 62%) but far lower precision (13% vs. 30%), leading to a weaker F1 score (17% vs. 40%).

**Figure 8:**
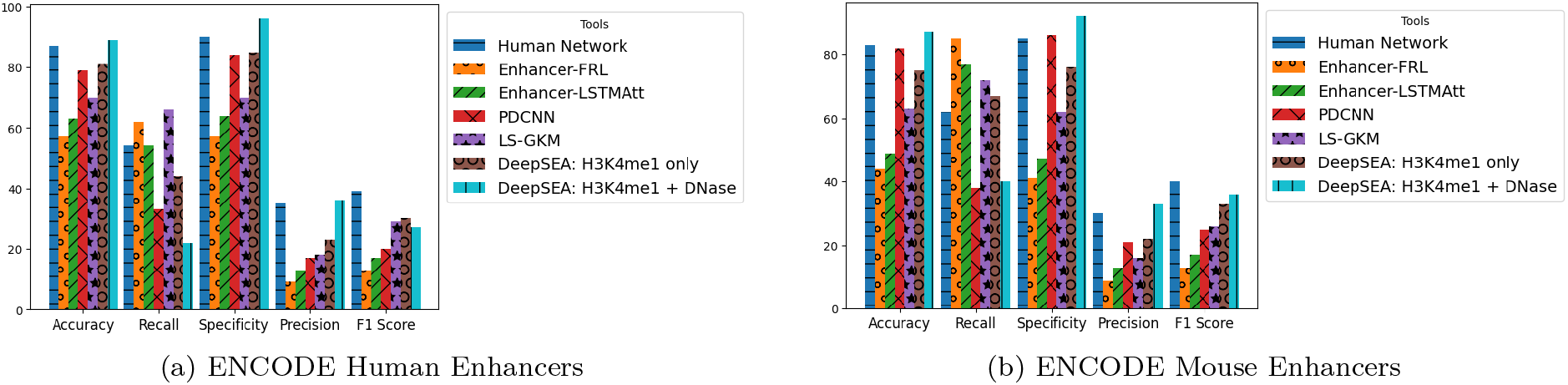
Performance comparison of enhancer classification tools on human and mouse ENCODE datasets. Bar plots show accuracy, recall, specificity, precision, and F1 score for six enhancer classification tools: our human network, Enhancer-FRL, Enhancer-LSTMAtt, PDCNN, LS-GKM, and two versions of DeepSEA, which is a sequence-to-expression tool. All tools were evaluated on the same test set for mouse enhancers. All tools except DeepSEA were evaluated on the same test set for the human enhancers. To evaluate DeepSEA, the human data were downsampled (40,000 enhancers and 400,000 controls) due to file size limits and the need to upload many files manually to DeepSEA’s website. Our human network consistently achieved the highest F1 score while maintaining competitive accuracy, recall, specificity, and precision.

When compared to PDCNN, our network again performed better across metrics. On the human data, our model achieved 87% accuracy and 42% F1, compared to PDCNN’s 79% accuracy and 20% F1. On the mouse data, PDCNN reached 82% accuracy and 25% F1, while our model matched its accuracy (83%) but achieved much stronger recall (62% vs. 38%) and F1 (40% vs. 25%).

In sum, our human network consistently outperforms Enhancer-FRL, Enhancer-LSTMAtt, PDCNN, and LS-GKM. It achieves superior F1 score on both the human and mouse datasets, providing the best balance between recall and precision. Despite being trained solely on human enhancers, our model generalizes strongly across species and experimental assays, including from snATAC-seq-derived CATlas enhancers to DNase I hypersensitivity-based ENCODE datasets, making it a reliable enhancer prediction tool.

### 2.17 *EnhancerDetector* Provides a Direct Enhancer-Classification Advantage over Sequence-to-Expression Models

Using the same ENCODE human and mouse enhancer datasets described in Section 2.16, we next compared *EnhancerDetector* to two general sequence-to-expression models, DeepSEA (previously evaluated on the CATlas human testing set under Section 2.15) and Enformer. These two tools predict chromatin features rather than enhancer labels directly. To ensure fairness, we adapted our evaluation to each model’s input requirements while maintaining the same enhancer-to-control ratios as in our other benchmarks.

DeepSEA requires 2000 bp inputs, so each 400 bp sequence from the ENCODE human and mouse test sets was symmetrically extended to 2000 bp. For consistency, we evaluated DeepSEA alongside the other enhancer-discovery tools using the same four cell-agnostic rules (H3K4me1; H3K4me1+DNase; H3K4me1+H3K27ac; H3K4me1+H3K4me3). We preserved the 1:10 enhancer-to-control ratio throughout. We evaluated it on the entire mouse dataset, but had to downsample the human data (40,000 enhancers and 400,000 controls) due to file size limitations and the manual effort required to upload a large number of files to DeepSEA’s website. Summary results are shown in Figure 8. On human data, DeepSEA achieved 81–89% accuracy and 15–30% F1 (depending on feature-rule choice), compared to *EnhancerDetector* ‘s 87% accuracy and 39% F1. On mouse data, DeepSEA reached 75–87% accuracy and 25–36% F1, while *EnhancerDetector* achieved 83% accuracy and 40% F1. Thus, given the 1:10 class imbalance where F1 is the most informative metric, *EnhancerDetector* consistently outperformed DeepSEA, even when their accuracy scores were similar.

To further benchmark *EnhancerDetector* against regulatory sequence models, we compared it to Enformer [44], a transformer-based deep learning model trained to predict thousands of functional genomic signals from long input sequences. Although Enformer is not explicitly designed as a binary enhancer classifier, it provides high-resolution predictions of chromatin accessibility and histone marks such as H3K4me1, H3K27ac, H3K4me3, and DNase hypersensitivity— features commonly associated with enhancers. We evaluated Enformer using our CATlas testing dataset. First, we identified transcription start sites (TSSs) where a 57,344 bp window on both sides (114,688 bp total) overlapped one or more annotated enhancer regions from our testing set. Matched control TSSs were selected with 114,688-bp-long windows. All selected sequences were expanded to 393,216 bp, as required by Enformer’s input length. We then selected 1,000 non-overlapping TSS-centered regions and ran Enformer on each of them. From the resulting output matrix, we extracted the predictions corresponding to any cell type for the four enhancer-associated features: H3K4me1, H3K27ac, H3K4me3, and DNase. For each enhancer or control region, we identified the bins in the output that overlapped the region and computed the mean signal for each of the four features. This resulted in four averaged columns per region (one for each feature). To normalize across regions, we computed the mean and standard deviation of each column and then transformed these values into z-scores by subtracting the column’s mean and dividing by the standard deviation. Using these normalized feature scores, we applied the same rule-based strategies as used for our DeepSEA assessment (H3K4me1 alone, H3K4me1 + H3K4me3, H3K4me1 + DNase and H3K4me1 + H3K27ac), for each rule, we computed both AUROC and AUPRC. Note that we did not compute traditional classification metrics (e.g., precision, recall at a threshold), as Enformer outputs continuous chromatin signal predictions rather than binary labels. Thus, interpreting a specific threshold as “active” is less meaningful and would introduce arbitrary cutoffs.

For *EnhancerDetector*, we used the same set of 114,688 bp TSS-centered regions and segmented them into overlapping 400 bp windows with a 25-bp stride. We then ran *EnhancerDetector* on each window and then took the windows that overlapped with the enhancer or control regions. We then calculated the average for those windows; from these average scores, we calculated both AUROC and AUPRC for *EnhancerDetector*. This setup allowed us to directly compare Enformer’s inferred chromatin profiles to *EnhancerDetector* ‘s enhancer probabilities.

Using the normalized feature scores derived from Enformer’s chromatin predictions, we evaluated the performance of four feature-based enhancer rules across the CATlas testing set. It is important to note that due to the relative scarcity of enhancer regions overlapping with TSS-centered regions, the selected dataset was skewed toward controls, resulting in an approximate enhancer-to-control ratio of 1:7.5. As shown in Figure 9, the combination of H3K4me1 and DNase yielded the highest performance among the Enformer-derived rules, with an AUROC of 0.6927 and an AUPRC of 0.2064. This was followed closely by the H3K4me1-only rule (AUROC = 0.6847, AUPRC = 0.2038). The lowest performance was observed for the H3K4me1 + H3K27ac rule (AUROC = 0.6759, AUPRC = 0.1919). In contrast, *EnhancerDetector* directly predicted enhancer probability scores across the same regions and achieved an AUROC of 0.7326 and an AUPRC of 0.2769, outperforming all Enformer-based rules. These results highlight that while Enformer provides rich chromatin signal predictions that could infer enhancer activity, it does not offer direct enhancer classification. Users must infer activity by combining multiple features and applying custom thresholds. In contrast, *EnhancerDetector* simplifies this process by directly predicting enhancer status from 400 bp sequences, making it a more streamlined and user friendly option for enhancer discovery.

**Figure 9:**
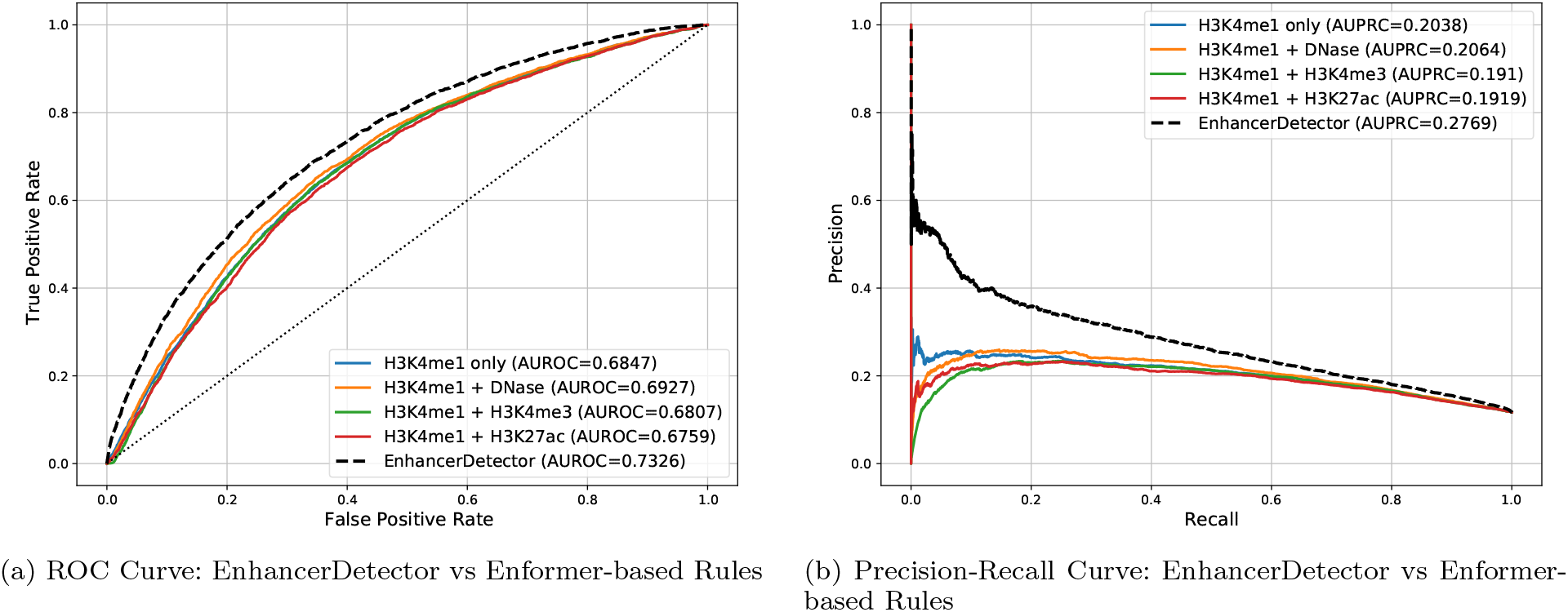
Comparison of enhancer classification performance between *EnhancerDetector* and four feature-combination rules derived from Enformer’s predictions (H3K4me1, H3K4me1 + DNase, H3K4me1 + H3K4me3, and H3K4me1 + H3K27ac). Panel (a) shows the receiver operating characteristic (ROC) curves and area under the ROC curve (AUROC) values; panel (b) shows the precision-recall curves (PRC) and area under the precision-recall curve (AUPRC) values. *EnhancerDetector* consistently outperforms all Enformer-based rule combinations in both AUROC and AUPRC.

We further evaluated both models using our ENCODE mouse testing set, following the same procedures as with the human dataset. However, due to the smaller number of mouse enhancers and controls, we increased the number of control TSS-centered sequences from 1,000 to nearly 5,000. This adjustment yielded a more realistic evaluation, producing an enhancer-to-control ratio of approximately 1:7.4 closely matching the 1:7.5 ratio used in the human testing. Enformer was evaluated using its mouse model and features, while *EnhancerDetector* was tested using its fine-tuned mouse model. *EnhancerDetector* achieved comparable results to Enformer with an AUROC of 0.8458. The best AUROC was obtained with the H3K4me1 + DNase combination (AUROC = 0.8621), followed by H3K4me1 + H3K27ac (0.8556), H3K4me1 + H3K4me3 (0.8548), and H3K4me1 alone (0.8540). In terms of AUPRC *EnhancerDetector* also showed comparable results to all Enformer based rules. The best AUPRC among the Enformer rules was 0.4038 (H3K4me1 + DNase), compared to 0.4256 due to *EnhancerDetector*. The results stayed comparable with 0.3967, 0.3945 and 0.3915 from H3K4me1 + H3K27ac, H3K4me1 only and H3K4me1 + H3K4me3, respectively. These results demonstrate that *EnhancerDetector* is able to match Enformer derived enhancer rules in both AU-ROC and AUPRC for mouse sequences, while offering a more direct and streamlined solution for enhancer prediction without the need for handcrafted feature combinations or post-hoc rule design.

Overall, *EnhancerDetector* delivers higher or comparable performance while offering a critical practical advantage: DeepSEA and Enformer require users to select among multiple chromatin-mark rules or thresholds, with the “best” choice varying by dataset. In contrast, *EnhancerDetector* outputs a direct enhancer probability from short 400-bp-long sequences, eliminating post hoc rule-tuning and providing a simpler, more consistent solution for enhancer discovery across species and datasets.

## 3 Discussion

### First-pass regulatory annotation in genomes

The rapid growth of genome sequencing has created a widening gap between the availability of high-quality genome assemblies and the functional annotation of their non-coding regulatory elements. Enhancers are particularly difficult to annotate because their activity is often cell-type-specific, their sequence conservation can be limited, and experimental assays remain unavailable for most species. In this study, we developed *EnhancerDetector*, a sequence-based framework for identifying putative enhancers directly from DNA sequence. The results support the idea that enhancer sequences contain recurring features that can be learned from diverse training data and transferred across species and experimental assays. Importantly, *EnhancerDetector* does not require chromatin marks, expression profiles, or cell-type-specific experimental data at prediction time, making it well suited for first-pass regulatory annotation in genomes for which functional genomic resources are sparse or absent.

### *EnhancerDetector* as an adaptable frame-work

A practical application of *EnhancerDetector* is the first-stage annotation of newly sequenced genomes. In such a workflow, a newly assembled genome could first be screened using *EnhancerDetector* to identify genomic windows with high enhancer probability scores. These candidate enhancers could then be prioritized according to score, conservation, or proximity to genes of interest. When a modest number of enhancer annotations becomes available for the target species, the human-trained model can be fine-tuned to improve species-specific performance. Our fine-tuning experiments indicate that a modest set of experimentally validated enhancers can markedly enhance performance, highlighting *EnhancerDetector* as a flexible framework instead of a one-size-fits-all classifier.

### Assigning cell types to putative enhancers

The predictions produced by *EnhancerDetector* can also be integrated with complementary tools that address different aspects of enhancer annotation. For example, after *EnhancerDetector* identifies putative enhancers in a newly sequenced genome, *EnhancerMatcher* can be used to compare these candidates with a small number of known cell-type-specific enhancers and infer likely cell-type activity using as few as two confirmed examples [65]. Together, these tools form a two-stage annotation strategy: *EnhancerDetector* first identifies candidate enhancer sequences across the genome, and *EnhancerMatcher* then classifies those candidates according to likely cell-type activity. This workflow is particularly useful for newly sequenced genomes.

### Systematic *in silico* experimentation

In addition to enhancer prediction, *EnhancerDetector* offers an interpretable framework for investigating the sequence grammar underlying enhancer activity. Class activation maps identify the regions of an input sequence that contribute most strongly to enhancer classification. These maps can be used not only for visualization but also as guides for systematic *in silico* experiments. In this study, perturbing CAM-identified regions reduced enhancer classification more strongly than perturbing other regions, while inserting high-scoring regions into non-enhancer sequences increased enhancer predictions. These experiments suggest that CAMs can help identify candidate sequence elements that contribute to enhancer identity. More broadly, CAM-guided perturbation can be used to test hypotheses about motif spacing, motif orientation, local sequence context, motif density, and the contribution of flanking sequence to enhancer activity.

### Interpreting experimental validation

The experimental validation in transgenic flies provides additional support for the utility of *EnhancerDetector* in prioritizing enhancer candidates. Five of six tested sequences drove reporter expression, and four produced patterns consistent with prior knowledge of the enclosing regulatory regions or likely target genes. These results suggest that *EnhancerDetector* can identify biologically active enhancer fragments within longer regulatory sequences. At the same time, the validation experiments illustrate the complexity of enhancer biology: some activities were partial, context-dependent, or observed outside the expected tissue or developmental context. Therefore, *EnhancerDetector* predictions should be interpreted as prioritizations of putative enhancer activity rather than definitive functional annotations. This distinction is especially important when applying the method to newly sequenced genomes, where experimental validation and biological context may be limited.

### Limitation of the positive datasets

An important limitation is that the positive datasets used for training and evaluation are enriched for enhancers but do not consist exclusively of experimentally confirmed enhancers. Genome-scale enhancer prediction is inherently challenging because available enhancer annotations are usually inferred from chromatin accessibility, transcriptional activity, or curated regulatory evidence, all of which provide incomplete and context-dependent views of enhancer function. We therefore interpret these datasets as high-confidence collections of putative enhancers rather than complete ground-truth enhancer catalogs. Despite this limitation, they represent the most comprehensive enhancer resources currently available across human, mouse, and fly, and they provide a practical basis for training and evaluating sequence-based enhancer discovery models.

### Control datasets

An important consideration in sequence-based enhancer prediction is whether the model is learning enhancer-associated regulatory features rather than simpler sequence-composition biases. We addressed this concern by evaluating *EnhancerDetector* against five distinct control datasets designed to test potential confounders, including sequence length, GC content, repetitive elements, and k-mer composition. Specifically, we compared enhancers with length-matched controls with and without repeats, length- and GC-matched controls with and without repeats, and shuffled enhancer-derived sequences that preserve local sequence composition. Performance across these increasingly stringent control settings indicates that the classifier is not driven solely by simple differences in length, GC content, repeat content, or k-mer composition. Although no computational control set can eliminate all possible sources of bias, these complementary negative datasets provide a strong framework for evaluating enhancer prediction under challenging background assumptions.

In summary, *EnhancerDetector* provides a practical and interpretable framework for enhancer discovery from DNA sequence alone. Its ability to generalize across species, experimental assays, and control settings supports its use as a first-stage tool for prioritizing putative enhancers in newly sequenced genomes. The fine-tuning strategy further allows the model to be adapted as species-specific enhancer annotations become available, while CAM-based interpretation enables systematic *in silico* experiments to investigate enhancer sequence grammar. When combined with *EnhancerMatcher*, putative enhancers predicted by *EnhancerDetector* can be further classified according to likely cell-type activity. Together, these features make *EnhancerDetector* a useful component of scalable regulatory genome annotation workflows and a platform for generating experimentally testable hypotheses about enhancer-associated sequence features.

## 4 Conclusion

Understanding how enhancers encode regulatory information in DNA is a central challenge in genomics, particularly as genome sequencing continues to advance more rapidly than functional annotation. A key open question is whether enhancers share an intrinsic, sequence-encoded identity—here termed enhancerness—that distinguishes them from other genomic regions independently of species, cell type, or experimental assay. In this context, establishing the existence and learnability of enhancerness is both a fundamental biological problem and a practical requirement for scalable genome annotation. Below, we summarize how our results address this challenge and demonstrate the implications of learning enhancer identity directly from DNA sequence.

In this study, we introduced *EnhancerDetector*, a convolutional neural network-based framework for accurate and interpretable identification of transcriptional enhancers across cell types and across species. *EnhancerDetector* consistently outperforms existing enhancer prediction methods, achieving superior F1 scores on both human and mouse datasets. The tool maintains an excellent balance between recall and precision, which are critical for genome-wide applications. *EnhancerDetector* directly classifies enhancers from DNA sequences, outperforming or achieving comparable performance to indirect sequence-to-expression models like DeepSEA and Enformer, which require additional thresholding and interpretation of chromatin feature predictions.

Our model generalizes well to independent datasets derived from diverse experimental assays, including snATAC-seq, CAGE, and DNase-seq.

A key strength of *EnhancerDetector* is its cross-species transferability: the model trained on human enhancers performs robustly on mouse and fly data and can be effectively fine-tuned using as few as 20,000 enhancers—a relatively small dataset in genomic studies. This capability makes it particularly valuable for enhancer discovery in newly sequenced genomes with limited experimental data.

We harnessed model interpretability through class activation maps to highlight sequence regions most predictive of enhancer activity. Computational perturbation analyses—including knockout, insertion, and sequence context disruption—confirmed the biological relevance of these regions. These analyses revealed that enhancers generally possess a core functional region supported by a broader sequence context. Our *in silico* experiments confirmed the directional nature of enhancers.

For experimental validation, we tested six putative enhancers in transgenic flies. Five (83%) drove reporter gene expression, and four (67%) showed expression patterns consistent with prior literature or their putative target genes.

In sum, *EnhancerDetector* offers a high-performing, biologically grounded, and easily adaptable platform for enhancer discovery. It lays a strong foundation for next-generation computational regulatory genomics, enabling accurate enhancer identification at scale—a critical capability as genomic data expand exponentially.

## Declarations

Additional files

Additional file 1: BLAST output of true positive mouse enhancers versus the human genome. File format: text (.txt).

## List of abbreviations

AI: Artificial Intelligence
AUPRC: Area Under the Precision-Recall Curve
AUROC: Area Under the Receiver Operating Characteristic Curve
bp: Base Pair
CAM: Class Activation Map
CNN: Convolutional Neural Network
eRNA: Enhancer RNA
GC: Guanine-Cytosine
GFP: Green Fluorescent Protein
gkm-SVM: Gapped k-mer Support Vector Machine
H3K4me1: Histone H3 Lysine 4 Monomethylation
H3K4me3: Histone H3 Lysine 4 Trimethylation
H3K27ac: Histone H3 Lysine 27 Acetylation
KL: Kullback-Leibler
LGR: Length GC Repeat
LGNR: Length GC No Repeat
LNR: Length No Repeat
LR: Length Repeat
LS-GKM: Large-Scale Gapped k-mer Support Vector Machine
PRC: Precision-Recall Curve
RNN: Recurrent Neural Network
ROC: Receiver Operating Characteristic
SVM: Support Vector Machine
TFBS: Transcription Factor Binding Site
TSS: Transcription Start Site

## Acknowledgements

We thank Jack Leatherbarrow and James Weiser for their assistance with the experimental validations.

## Availability of data and materials

The *EnhancerDetector* code is available at https://github.com/BioinformaticsToolsmith/EnhancerDetector and archived at https://doi.org/10.5281/zenodo.15531292. The software is available under the Affero General Public License version 3 for academic use; commercial use requires an alternative license.

## Authors’ contributions

L.M.S. developed the software, analyzed the results, and wrote the manuscript. G.S.-L. performed the in vivo validation experiments. M.S.H. conceived the original idea, analyzed the results, interpreted the in vivo validation experiments, and wrote and reviewed the manuscript. H.Z.G. conceived the original idea, supervised the project, developed the software, analyzed the results, and wrote and reviewed the manuscript. All authors read and approved the final manuscript.

## Ethics approval and consent to participate

Not applicable.

## Consent for publication

Not applicable.

## Author details

